# Cardiovascular disease risk factors induce mesenchymal features and senescence in cardiac endothelial cells

**DOI:** 10.1101/2020.10.21.349472

**Authors:** Karthik Amudhala Hemanthakumar, Fang Shentong, Andrey Anisimov, Mikko I. Mäyränpää, Eero Mervaala, Riikka Kivelä

**Affiliations:** Wihuri Research Institute, Helsinki, Finland; Stem cells and Metabolism Research Program, Research Programs Unit, Faculty of Medicine, University of Helsinki, Helsinki, Finland; Translational Cancer Medicine Research Program, Research Programs Unit, Faculty of Medicine, University of Helsinki, Helsinki, Finland; Pathology, Helsinki University and Helsinki University Hospital, Helsinki, Finland; Department of Pharmacology, Faculty of Medicine, University of Helsinki, Helsinki, Finland

**Keywords:** Aging, Obesity, Exercise, Endothelium, Cellular reprogramming, Heart, TGF-β

## Abstract

Aging, obesity, hypertension and physical inactivity are major risk factors for endothelial dysfunction and cardiovascular disease (CVD). We applied fluorescence-activated cell sorting (FACS), RNA sequencing and bioinformatic methods to investigate the common effects of CVD risk factors on cardiac endothelial cells (ECs). Aging, obesity and pressure overload all upregulated pathways related to TGF-β signaling and mesenchymal gene expression, inflammation, vascular permeability, oxidative stress, collagen synthesis and cellular senescence, whereas exercise training downregulated most of the same pathways. We identified collagen chaperone SerpinH1/HSP47 to be significantly increased by aging and obesity and repressed by exercise training. Mechanistic studies demonstrated that SERPINH1/HSP47 in human ECs changed cell morphology and increased mesenchymal gene expression, while its silencing inhibited collagen deposition. Our data demonstrate that CVD risk factors significantly remodel the transcriptomic landscape of cardiac ECs to acquire senescence and mesenchymal features. SERPINH1/HSP47 was identified as a potential therapeutic target in ECs.

## Introduction

According to WHO, cardiovascular diseases (CVD) account for 10% of the global disease burden and constitute the number one cause of death in the Western world. CVD are mainly caused by behavioral (physical inactivity, unhealthy diet) and metabolic (obesity, hypertension, diabetes, high cholesterol) risk factors ^1^. Aging, however, is by far the biggest contributor to CVD, and aging population is becoming an enormous challenge worldwide.

The heart contains a dense vascular network, and endothelial cells (EC) are indeed the most abundant cell population in the adult mouse heart ^2^. In addition to their transport function, ECs are defined to control vasomotor tone, maintain vascular homeostasis, regulate angiogenesis, and to establish bidirectional communication with other cell types and organs via paracrine signaling mechanisms ^3–7^. ECs are found to be highly adaptive to physiological stimuli during normal growth and development ^8, 9^, and the diversity of ECs in different tissues has now been acknowledged. ECs are also maladaptive to a spectrum of pathological events involving e.g. inflammation or oxidative stress ^10, 11^, and the development of heart diseases is strongly linked to endothelial dysfunction and impaired vascular remodeling. However, the molecular cues, which cause maladaptation and dysfunction of ECs in the heart in response to pathological signals, remain elusive.

Physical inactivity increases the incidence of several chronic diseases, whereas regular exercise training has positive effects on most of our tissues ^12^. Because microcirculation is present in every organ in the body, ECs have a unique ability to influence the homeostasis and function of different tissues, and they are potentially a major cell type mediating the positive effects of exercise throughout the body. Although the cardiac benefits of exercise are clear and there have been major advances in unraveling the molecular mechanisms, the understanding of how the molecular effects are linked to health benefits is still lacking^12^. Especially, the effects on ECs have not been characterized.

We hypothesized that the major CVD risk factors aging, obesity and pressure overload will induce adverse remodeling of cardiac EC transcriptome ^11, 28, 29^, whereas exercise training would provide beneficial effects ^8, 9^. Both physiological and pathological stimuli significantly modified the cardiac EC transcriptome. Intriguinly, our results demonstrated that CVD risk factors promoted activation of transforming growth factor-β (TGF-β) signaling, cellular senecence and induced mesenchymal gene expression in cardiac EC, whereas exercise training promoted opposite protective effects.

## Results

### Exercise training and CVD risk factors modulate cardiac EC number, vascular density and transcriptome

To mimic the effect of the most common CVD risk factors (aging, obesity, pressure overload/hypertension and physical inactivity), we used adult C57BL/6J wild type mice in the following experimental groups: aged (18 months) vs. young (2 months) mice, high-fat diet induced obesity (14 weeks HFD) vs. lean mice, transverse aortic constriction (TAC) vs. sham-operated mice and exercise training (progressive treadmill running for 6 weeks) vs. sedentary mice (**Figure S1A-B**). Exercise trained mice showed improved ejection fraction compared to the sedentary mice, whereas aging, HFD and TAC resulted in impaired heart function (**Figure S1C-F and Supplementary Table 1**). HFD also induced marked weight gain, increased fat mass and impaired glucose tolerance (**Figure S1G-I**). Left ventricular (LV) mass was increased in aged, HFD-treated and TAC mice (**Supplementary Table 1**). Exercise training also slightly increased LV mass, which reflects mild physiological hypertrophy often observed in endurance-trained athletes ^30^ (**Supplementary Table 1**).

Exercise training significantly increased, whereas aging, HFD and TAC decreased the percentage, count and mean fluorescence intensity of the cardiac ECs (CD31^+^CD140a^-^ CD45^-^Ter119^-^DAPI^-^) compared to the controls, when analyzed by FACS (**Figure 1A-B, Figure S2A-D**). This was also demonstrated by immunohistochemistry for CD31-positive coronary vessels (**Figure 1C-D**). The cardiac ECs were gated and sorted by FACS (Figure S3A), and the isolated ECs were first analyzed by quantitative PCR analysis, which indicated significant enrichment of EC markers Cdh5 and Tie1 in the sorted fraction compared to whole heart or other cardiac mononuclear cells (**Figure S3B**). In addition, isolation resulted in 87.4±1.9% cell viability and RNA purification strategy yielded intact and stable RNA with average RNA integrity number (RIN) of 8.7 (Figure S3C-D). RNA sequencing of isolated ECs was used to profile the expression pattern of cardiac EC transcripts in different experimental groups. Two-dimensional principal component analysis of the EC transcriptomes exhibited significant proportion of variance in the gene expression pattern, which can be attributed to the treatment-induced changes in cardiac EC transcriptome (**Figure S4A-E**). Notably, unsupervised hierarchical clustering of EC data sets for all experimental interventions (sedentary, exercise trained, young, aged, sham, TAC) revealed consistent clustering and high degree of similarity in the gene expression pattern (**Figure S4F-J**). The analysis for differentially expressed genes (DEGs) showed a large number of up- and downregulated genes especially in aged, obese and TAC-operated mice followed by a smaller number of affected genes in exercise trained mice. The number of significantly up- and downregulated genes with the false discovery rate 0.05 for each treatment are shown in the MA plots and the top 50 DEGs for each treatment are presented by heat maps (**Figure 2A-E, F-J**).

**Figure 1.**
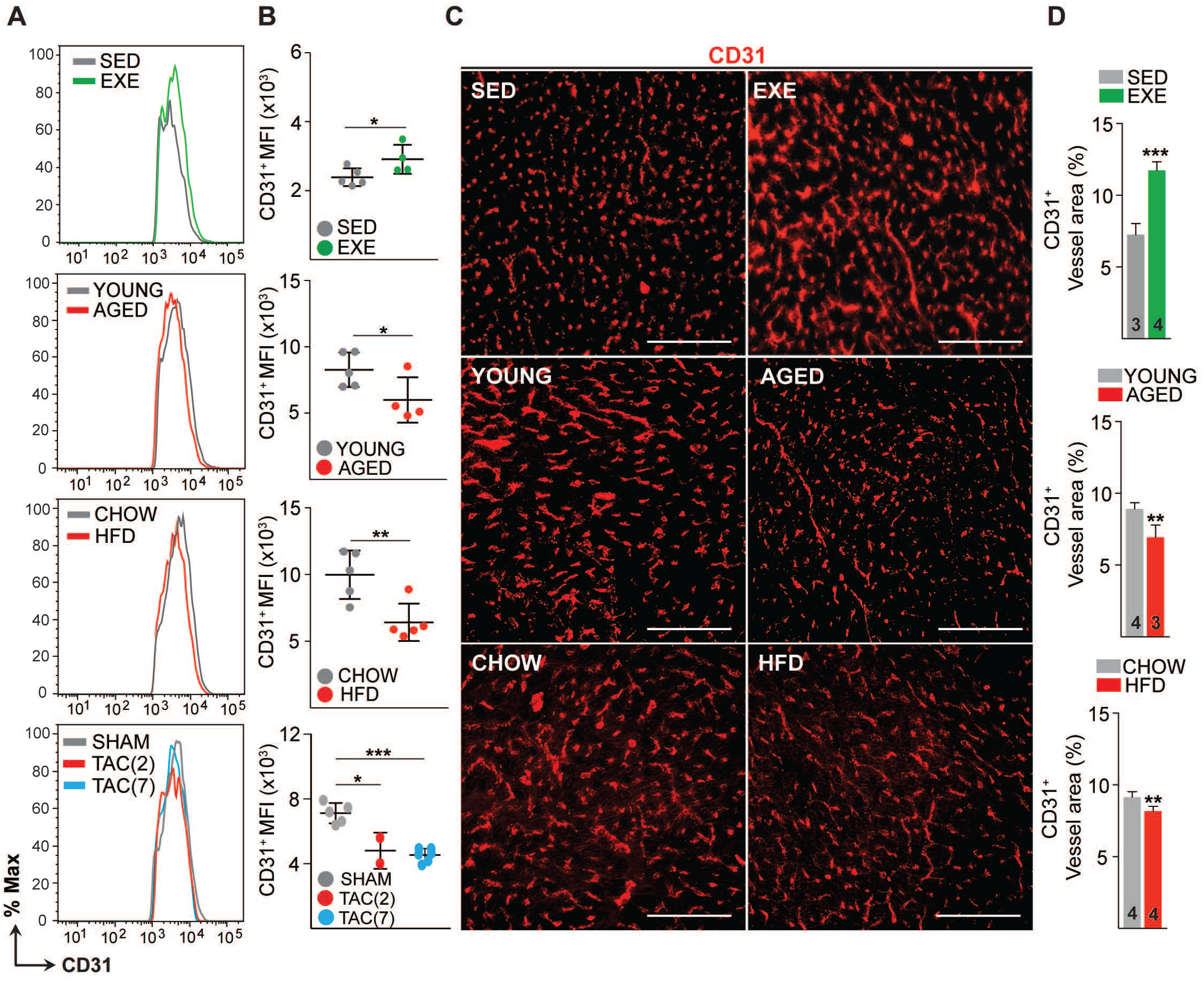
Effects of exercise training, aging, obesity and pressure overload on cardiac endothelial cell number and vascular density. A-B. FACS analysis and quantification of mean fluorescence intensity (MFI) of the cardiac endothelial cells (CD31+CD140a-CD45-Ter119-DAPI-) in various mice models. **C-D.** Representative immunofluorescence images and quantification of CD31+ blood vessel area (%) in the heart. Scale bar, 100μm. Data is presented as mean ± SEM. Student’s t test was used, *p<0.05, **p<0.01, ***p<0.001 (N=3-5 mice/group). In the panel **B**, each color-coded circle (Red, Green and Black) indicates an individual biological sample. In panel **D**, number of mice in each experimental group are indicated in the respective graph, N=3-5 mice/group.

**Figure 2.**
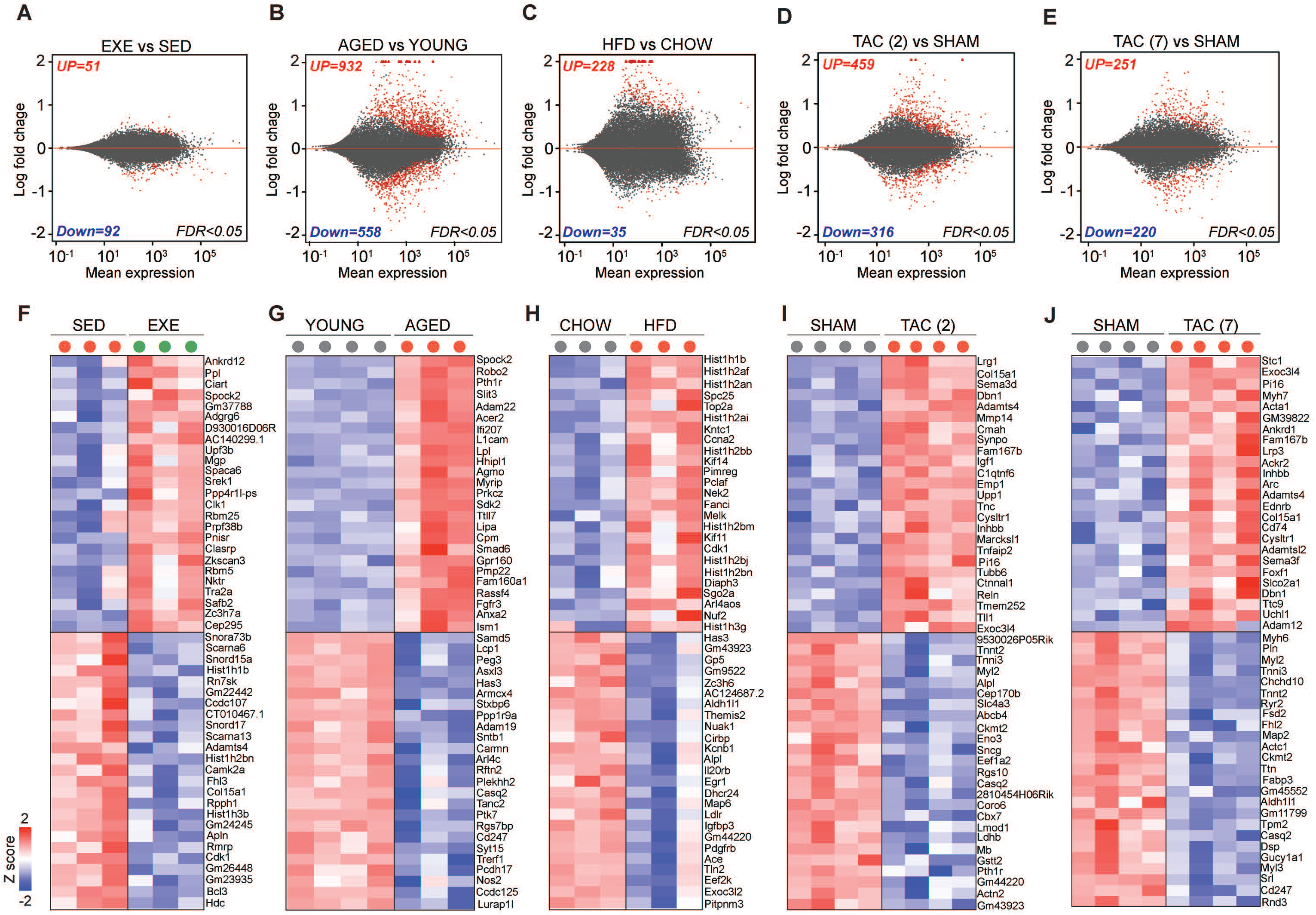
Transcriptomic changes in cardiac endothelial cells from exercise trained, aged, obese and TAC-treated mice. A-E. MA (log ratio over mean) plots showing the number of differentially expressed genes in cardiac ECs for each experiment. Number of significantly up- and downregulated genes with the FDR (Benjamini-Hochberg adjusted pvalue) threshold of 0.05 are indicated in the plots. **F-J.** Top 50 differentially expressed genes in cardiac ECs of the indicated experimental groups. In the heatmap, each color-coded circle (Red, Green and Black) indicates an individual biological sample within each experimental group. N=3-4 mice/group.

### CVD risk factors induce senescence and TGF-β signaling together with mesenchymal gene expression in cardiac ECs

To understand biological functions of the differentially expressed genes (DEG), we used PANTHER classification analysis (**Figure 3A**). The analysis revealed that genes related to EC development, adherence junction organization, IGFR signaling, adrenomedullin receptor signaling, and mitochondria were upregulated by exercise training. Furthermore, exercise training downregulated pathways related to cellular aging, vascular membrane permeability, negative regulation of angiogenesis, TGF-β1 production, collagen activated tyrosine kinase signaling, and ossification. In contrast, pathways related to TGF**-**β, IFN*α*, TNF*α*, oxidative stress, EC differentiation, vascular permeability, cell aging, collagen synthesis, SMAD signaling and mesenchymal cell development were highly enriched in cardiac EC from both aged and obese mice. Downregulated pathways in these mice included tissue and lipid homeostasis, ECM assembly, tube morphogenesis, cell adhesion, cell number maintenance, EC proliferation, vasculature development, artery development, and NOTCH signaling. Pressure overload activated pathways such as cellular response to TGF-βR2 activation of fibrotic pathways, inactivation of cell survival pathways Erk1/2 and MAPK, and ossification process, whereas cellular homeostasis and vasculature development were repressed.

**Figure 3.**
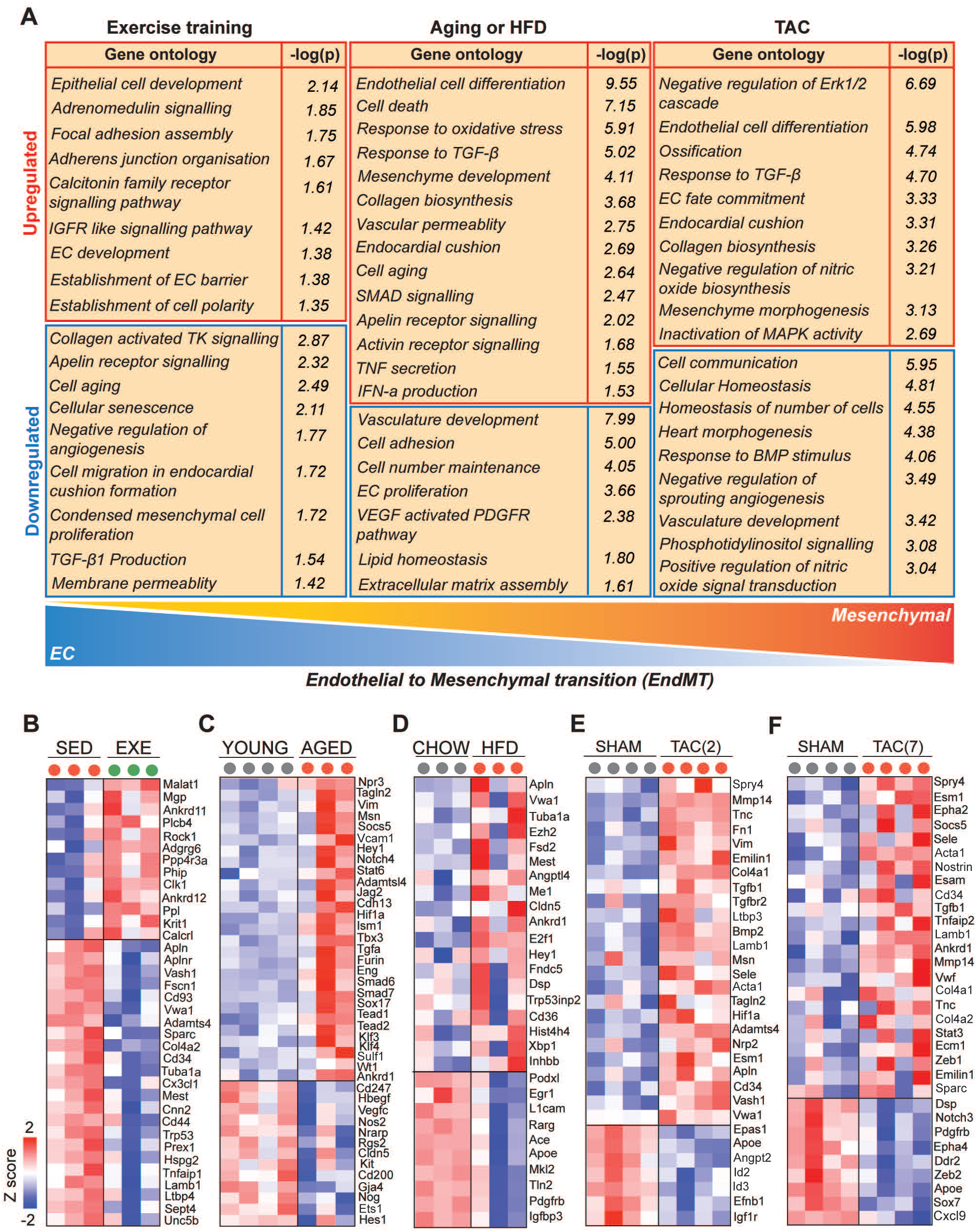
Cardiovascular disease risk factors activate mesenchymal gene expression in cardiac ECs. **A.** Gene ontology analysis of the up- and downregulated genes. Note the opposite changes by exercise training compared to the CVD risk factors. **B-F.** Heatmaps showing the differential gene expression of endothelial and mesenchymal genes previously associated to endothelial-to-mesenchymal transition (EndMT). Genes are selected based on published datasets (references are found in the supplementary table 5A-E). In the figure panel A-F, the up- and downregulated genes with the FDR (Benjamini-Hochberg adjusted pvalue) threshold of 0.05 were considered. In the heatmap, each color-coded circle (Red, Green and Black) indicates an individual biological sample within each experimental group. N=3-4 mice/group.

Comparison of the GO biological terms, which were significantly affected by exercise training and the CVD risk factors, demonstrated clear opposite effects on the EC transcriptome. Aging and HFD promoted oxidative stress response, activation of inflammatory and fibrosis pathways and cellular aging, and inhibited pathways regulating cell number maintenance, proliferation and lipid homeostasis. Exercise training, in turn, promoted EC homeostasis and vascular growth, and prevented vascular aging, inflammation and pathological activation.

Because the gene ontology categories indicated upregulation of genes and pathways associated to mesenchymal development and endothelial-to-mesenchymal transition (EndMT) by CVD risk factors, we reviewed our differentially expressed gene sets for the expression of selected endothelial and mesenchymal markers based on the previously published data sets (**Supplementary Table 5A-E**). We found significant upregulation of many mesenchymal markers and downregulation of EC genes in aged and obese mice (**Figure 3C-D**). After two weeks of TAC, we also observed upregulation of several mesenchymal markers, whereas after seven weeks of TAC, there was both up- and downregulation of the EC and mesenchymal markers, indicating possible reversal of the process (**Figure 3E-F**). Strikingly, exercise training downregulated several EndMT genes (Fscn1, Cd93, Vwa1, Sparc, Tuba1a, Cd44, Trp53, Col4a2, Mest, Cnn2, Tnfaip1, Lamb1, Ltbp4, Unc5b), the angiogenesis inhibitor gene Vash1, and the endothelial activation marker Apln and its receptor Aplnr, whereas it upregulated the expression of Malat1, Mgp, Krit1 and Calcrl (**Figure 3B**). We validated the results using an expanded set of samples by qPCR for Apln, Vim, Tgfbr2, Vash1, Sparc and Tgfb1 (**Figure S6A-F**).

### SerpinH1 expression is increased by aging and obesity and repressed by exercise training

To identify genes, which could mediate the negative effect of aging and obesity and the protective effects of exercise, we performed gene overlap analysis of DEGs from these three experimental interventions. We found 4 genes significantly affected by all treatments, of which 2 genes (SerpinH1/Hsp47 and Vwa1) were upregulated by aging and HFD, and downregulated by exercise training. The other two genes (Mest and Fhl3) were upregulated by HFD and downregulated by exercise training and aging (**Figure 4A-C**). We performed an in silico secretome analysis to characterize the properties of the identified genes using MetaSecKB database (**Figure 4D**). Both SerpinH1 and Vwa1 contain a signal peptide for secretion, indicating they could act as angiocrines in autocrine and/or paracrine fashion.

**Figure 4.**
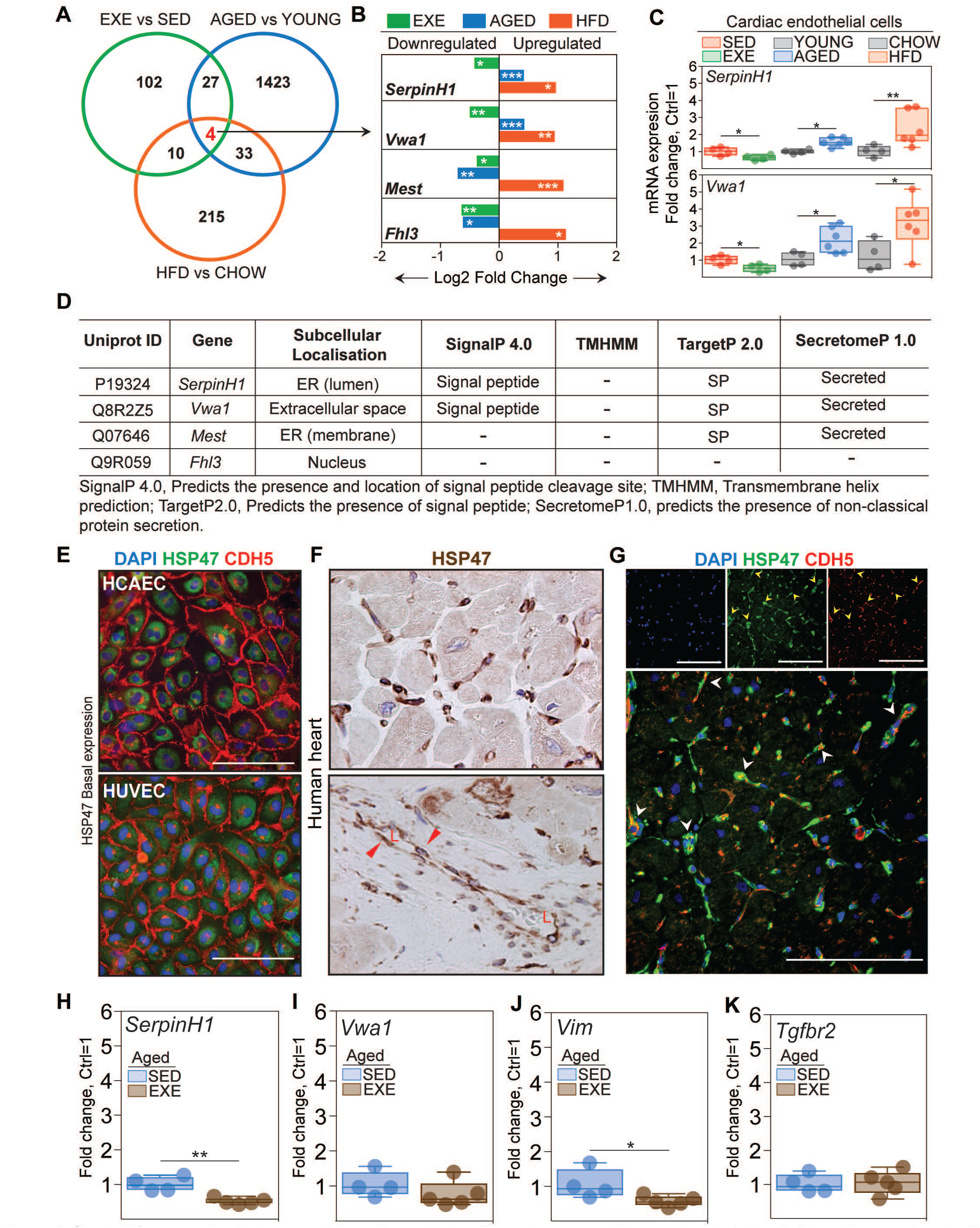
SerpinH1 expression is increased by aging and obesity and repressed by exercise training. **A.** Venn diagram showing the overlap of differentially expressed genes between the experiments. Four genes were identified to be significantly affected by aging, obesity and exercise (SerpinH1, Vwa1, Mest and Fhl3). **B.** Bar plot showing the expression pattern of these four genes. In panel **A** and **B**, the up- and downregulated genes with the FDR (Benjamini-Hochberg adjusted p-value) threshold of 0.05 were considered to be significant (N=3-4 mice/group). **C.** qPCR Validation of SerpinH1 and Vwa1 normalized to HPRT1 (N=4-6 mice/group). **D.** In silico secretome analysis of the identified genes. E-G. Representative immunofluorescent and immunohistochemistry images showing the expression of SERPINH1/HSP47 in human EC and human heart samples (red arrow head in the bottom panel F indicates the expression in large vessels and “L” indicate vessel lumen. White arrowheads in the panel G denote the co-expression of HSP47 and CDH5 in coronary vessels (yellow signal). **H-K.** mRNA expression of SerpinH1, Vwa1, Vim and Tgfbr2 in the cardiac ECs of sedentary and exercise trained aged mice (N=4-5/group). Scale bar 100 μm. Data is presented as mean ± SEM. Student’s t test was used, *p<0.05, **p<0.01, ***p<0.001.

We focused on SerpinH1/Hsp47, as it has a known role as a collagen chaperone and has been linked to fibrosis ^31^, making it an attractive candidate. We validated the endothelial SerpinH1/Hsp47 expression by qPCR (**Figure 4C**), and at single cell level using Tabula Muris database ^32^ and cardiac EC atlas from the Carmeliet lab ^33^. The scRNAseq analysis revealed that SerpinH1 is expressed in variety of cell types within the mouse heart, including fibroblasts, myofibroblasts, smooth muscle cells, endothelial cells, endocardial cells and to lesser extent in cardiomyocytes (**Figure S7A-D**). In ECs, SerpinH1 was found to be expressed throughout all endothelial cell clusters, with the highest expression in the apelin- high cluster marking activated ECs (**Figure S8A-F**). Interestingly, the expression of mesenchymal markers such as Tagln2, Vim and Smtn was also high in this cluster. Next, we analyzed the expression of SERPINH1/HSP47 in healthy human heart and in human cardiac ECs. Immunohistochemistry demonstrated SERPINH1/HSP47 to be highly expressed throughout the coronary vasculature and in fibroblasts in human heart, and weak staining was also detected in cardiomyocytes (**Figure 4E-G**, **Figure S7D**). In human cardiac ECs, HSP47 was localized perinuclearly, similarly to what has been demonstrated in other cells types, and consistent with the ER retention motif in its N-terminus (**Figure 4E**) ^34–36^. We also tested, if exercise training can attenuate the expression of SerpinH1, Vwa1 and selected markers of TGF-β signaling/EndMT also in aged mice. Of the studied genes, mRNA expression of SerpinH1 and Vimentin were significantly repressed by exercise and there was a tendency also for Vwa1 (**Figure 4H-K**).

### Overexpression of SERPINH1/HSP47 induces mesenchymal features in human ECs

To study the effects of SERPINH1/HSP47 in human ECs, we produced lentiviral vector encoding myc-tagged hSERPINH1/HSP47. Both human umbilical venous endothelial cells (HUVECs) and human cardiac arterial endothelial cells (HCAECs) were analyzed. HSP47 protein was localized similarly to the native protein (**Figure 5B**), and the overexpression was verified by western blotting (**Figure S9A**). Overexpression of HSP47 altered the cellular morphology characterized by impaired or discontinuous vascular endothelial cadherin junctions, increased stress fiber formation, and larger cell size (**Figure 5A-B**). Furthermore, analysis of EC and mesenchymal cell related transcripts demonstrated significant repression of EC markers (CD31, CDH5, TIE1, NRARP, ID1) and induction of a proliferation gene CCND1, and mesenchymal/EndMT markers (TAGLN, aSMA, CD44, VIM, NOTCH3, ZEB2, SLUG, FN1, VCAM1, ICAM1) (**Figure 5C**). VE-cadherin downregulation was also confirmed at protein level (**Figure 5D**) and increased aSMA expression by immunofluorescence staining.

**Figure 5.**
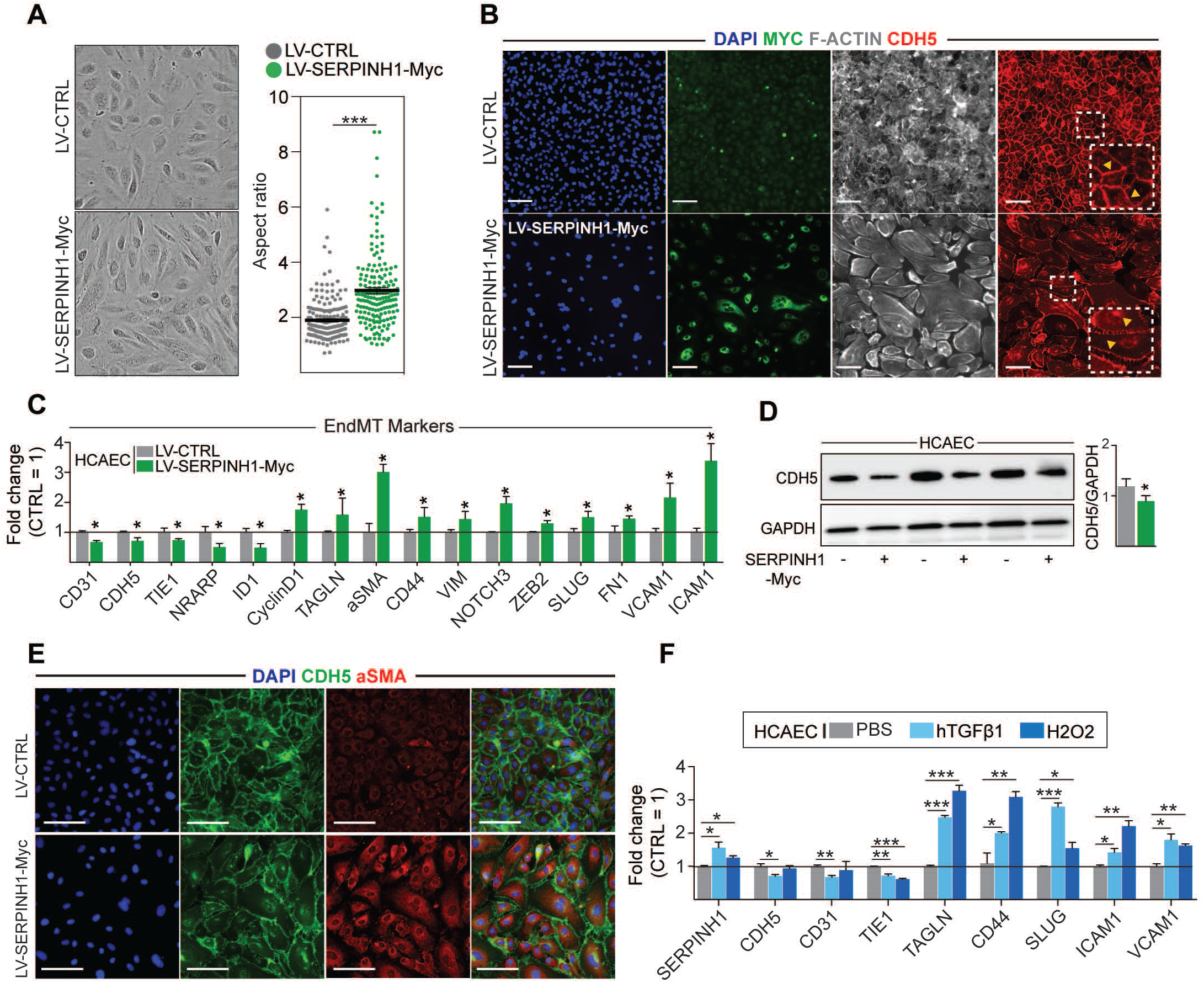
Overexpression of SERPINH1 modifies the EC phenotype and induces mesenchymal gene expression in human cardiac ECs. A. Representative phase-contrast images of live human cardiac arterial EC (HCAEC) transduced with LV-CTRL and LV-SERPINH1-Myc and quantification of the aspect ratio (length to width ratio) of the cell. **B.** Representative immunofluorescent images showing the expression of Myc-tagged SERPINH1 in green, F-Actin in grey and CDH5/VE-Cadherin in red. The insert within the white box shows magnified view of VE-Cadherin junctions in HCAECs. **C.** qPCR analysis of endothelial and mesenchymal markers in SERPINH1 overexpressing cells. **D.** Western blot analysis and quantification of CDH5/VE-cadherin expression in the SERPINH1 overexpressing HCAECs (normalized to GAPDH). **E.** Representative immunofluorescent images showing DAPI in blue, CDH5/VE-Cadherin in green and *α*-smooth muscle actin (aSMA) in red. **F.** qPCR analysis of SERPINH1 and EndMT markers in HCAECs stimulated with TGF-β1 (50ng/ml) or H2O2 for five days. In panel **A, C, D** and **F**, N=3 biological replicates/group were analyzed. Scale bar 100 μm. Data is presented as mean ± SEM. Student’s t test was used, *p<0.05, **p<0.01, ***p<0.001.

Transcriptomic changes pointed towards activated TGF**-**β signaling and oxidative stress in response to all of the CVD risk factors. Both are known to contribute to EC dysfunction and EndMT, thus we tested if they act as uptream regulators of HSP47. Indeed, our results show that TGF-β1 -treatment of HCAECs significantly upregulated the expression of SERPINH1 together with other known EndMT markers, and there was also small but significant induction of SERPINH1/HSP47 by hydrogen peroxide treatment (**Figure 5F**).

### SERPINH1/HSP47 is needed for collagen 1 deposition by ECs

To investigate the significance of SERPINH1/HSP47 depletion in human cardiac ECs, HCAECs were transduced with four independent shSERPINH1 lentiviral constructs. The constructs induced approximately 80% deletion of mRNA (**Figure 6D**). The cell morphology was not affected after two days (**Figure 6A**), but ten days of silencing significantly changed endothelial cell morphology and decreased the cell density in culture (**Figure 6B**), suggesting that SERPINH1/HSP47 might play a role in EC homeostasis and function. SERPINH1 silencing significantly inhibited collagen fibril deposition, detected by immunistochemistry for type 1 collagen (**Figure 6B, C**). Only the cells transduced with the construct #1 could produce some extracellular collagen 1, and these cells also survived better than the cells transduced with constructs #2, #3 or #4 (**Figure 6B, C**). Next, we treated the cells with TGF-β1 and hydrogenperoxide for five days to induce EndMT features, as described previously ^21, 37^. We used the shSERPINH1 (#1) construct, because from the other silencing constructs not enough cells survived for the experiments. The results indicated that silencing of SERPINH1 prevented the appearance of Taglin-positive cells, a commonly used readout for EndMT, which were observed in the control cells (**Figure 6E**). We also studied the effect of SERPINH1/HSP47 on cell proliferation/migration. In the scratch wound healing assay, overexpression of SERPINH1 significantly promoted wound closure (**Figure 7A, B**), whereas silencing of SERPINH1 for two days significantly decreased EC proliferation/migration. (**Figure 7C, D**).

**Figure 6.**
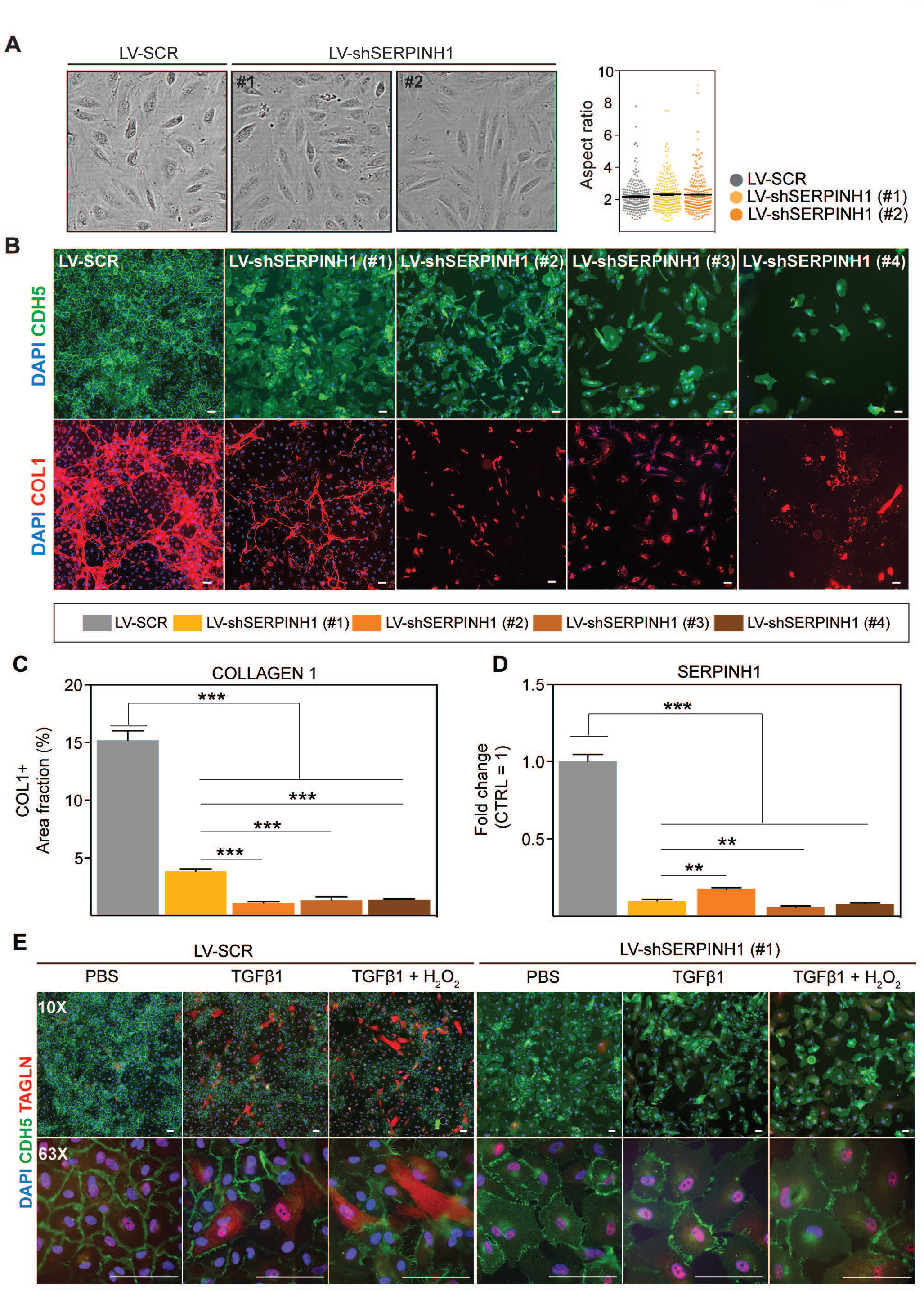
SERPINH1/HSP47 silencing in human cardiac EC inhibits collagen production and secretion. **A.** Representative phase contrast images of live HCAECs transduced with LV-SCR and LV-shSERPINH1 (#1 and #2) and quantification of the aspect ratio (length to width ratio) of the cells after two days of silencing. **B.** Representative CDH5/VE-Cadherin immunofluorescent images (green) showing the cell morphology and density after ten days of SERPINH1 silencing. Collagen 1 staining is shown in red, and quantification of Collagen 1 is shown in **C**. **D.** qPCR analysis of SERPINH1 deletion levels using four independent constructs. **E.** Representative immunofluorescent images showing TAGLN expression in the control and SERPINH1 silenced HCAECs treated with recombinant human TGF-*β*1 with and without H2O2 for five days. In the panel **A, C** and **D**, N=3 biological replicates/group were analyzed. Scale bar 100 μm. Data is presented as mean ± SEM. Student’s t test was used, *p<0.05, **p<0.01, ***p<0.001.

**Figure 7.**
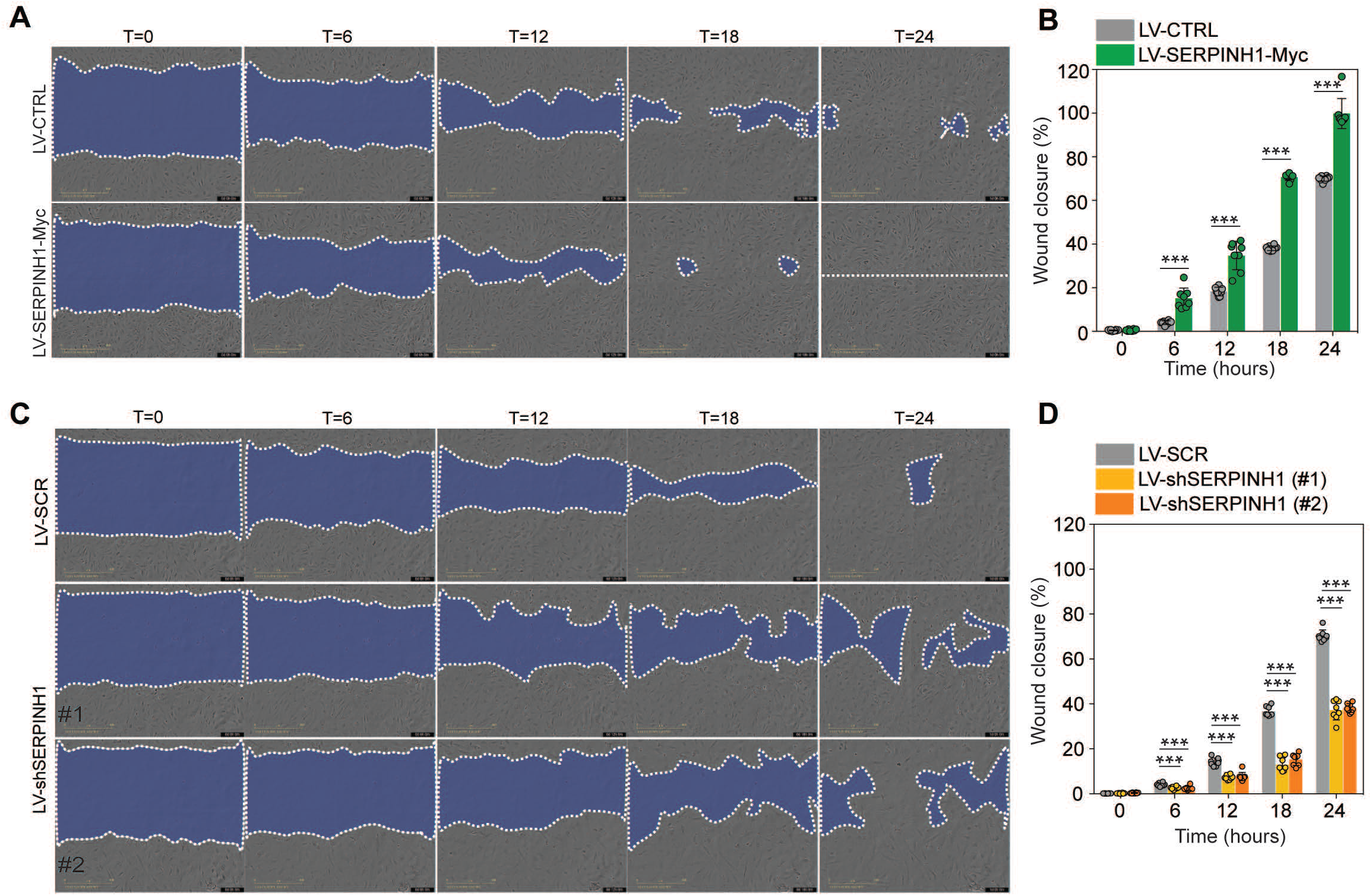
SERPINH1 overexpression enhances and silencing inhibits wound closure in vitro. A-B. Representative phase contrast images of scratch wound healing assay performed in HCAECs treated with LV-CTRL and LV-SERPINH1, and quantification of the wound closure (%) with respect to time (hours). **C-D.** Representative phase contrast images of scratch wound healing assay performed in HCAECs treated with LV-SCR and LV-shSERPINH1 (#1 and #2), and quantification of the wound closure (%) with respect to time (hours). In the panel A and C, the blue area within the white dotted region indicate the wound area. In the panel **B** and **D**, N=8 biological replicates/group were analyzed. Data is presented as mean ± SEM. Student’s t test was used, *p<0.05, **p<0.01, ***p<0.001.

## Discussion

Here we have used transcriptomic profiling to decipher how the major CVD risk factors aging, obesity and pressure overload remodel cardiac endothelial cells, and how the protective effects of exercise are mediated. The results demonstrate that the CVD risk factors activate transcriptional programs promoting cell aging, senescence, TGF-β activation, inflammation and oxidative stress in cardiac ECs. Importantly, exercise attenuated these same pathways, even in healthy mice. Furthermore, we found that aging, obesity and pressure overload induced mesenchymal gene programs in cardiac ECs, which can contribute to dysfunctional endothelium and CVD development. Analysis of potential disease-promoting genes identified SerpinH1/HSP47 to be induced by aging and obesity, while its expression was significantly repressed by exercise, even in old mice. Mechanistically, SERPINH1/HSP47 was induced by TGF-β and ROS, and the overexpression of SERPINH1/HSP47 increased cell size and stress fiber formation, weakened cell-cell junctions and promoted mesenchymal gene expression in human cardiac ECs. Immunohistochemistry of human hearts showed that HSP47 is abundantly expressed throughout the cardiac vasculature.

The largest dysregulation of the cardiac EC transcriptome was found in aged mice, followed by obesity and pressure overload. Exercise training affected a smaller number of transcripts, which can be accounted, at least partly, to the young and healthy control mice, which could move unrestrictedly in their home cages. Interestingly, however, most of the pathways activated by CVD risk factors were the same that were repressed by exercise training, highlighting the potential of physical activity to improve cardiovascular health via modulating endothelial cell phenotype and function. The positive effects of exercise on skeletal muscle and cardiac angiogenesis have been described previously ^38^, but exercise-induced molecular changes in ECs have not been characterized. Aging and obesity, on the other hand, are known to contribute to capillary rarefaction and/or dysfunction ^10, 11, 39, 40^, and another novel aspect in this study was the comparison of several CVD risk factors to identify common pathways and genes, which could drive the pathogenesis in cardiac disease, and could be considered as potential therapeutic targets. ECs would provide an attractive target for drug development, as they are the first cells to encounter drugs in the bloodstream.

Dysfunctional endothelium likely contributes to more diseases than any other tissue in the body as it affects all organs. On the other hand, endothelium could act as an important mediator of the health-promoting effects of exercise in a variety of tissues. Our finding that aging, obesity and pressure overload induce mesenchymal gene programs in cardiac ECs adds to the increasing evidence that activated endothelial TGF-β signaling and acquisition of mesenchymal features play an important role in the development of EC dysfunction and cardiac diseases ^13, 20, 41, 42^. Importantly, genes related to TGF-β production and cellular aging were repressed by exercise, highlighting the mechanisms behind the potential of exercise training in preventing and delaying the development of CVD. The activation of TGF-β signaling pathway has been implicated as a driving force for EndMT ^21, 23–26^. Several studies have recently suggested that EndMT could contribute to the development of various cardiovascular diseases ^16 13, 14, 27^, but currently there is a lack of understanding of the causal relationships and mechanisms linking EndMT and CVD ^13^. Furthermore, whether the transition from ECs to mesenchymal cells occurs completely in various CVDs is still actively debated in the literature. It has been suggested that pathological EC activation will result in acquired EndMT features e.g. expression of mesenchymal genes, without full transformation from one cell type to another ^44^. This is in line with our findings, as only cells with high CD31 expression and with no expression of CD45, CD140a and Ter119 were included in our analyses. Thus, all the analyzed cells were endothelial cells, but in the CVD risk factor groups they demonstrated increased mesenchymal marker expression. Long-term lineage tracing of ECs in response to CVD risk factors would provide further knowledge if and to what extent full transformation of ECs to mesenchymal cells occurs in cardiac vasculature. Our results, however, demonstrate that ECs acquire mesenchymal features due to CVD risk factors, which likely results in EC dysfunction even without EndMT.

To identify possible pathology-driving genes, which would be common for several risk factors, we performed gene overlap analysis using all data sets. Two genes, SerpinH1 and Vwa1, were found to be significantly increased by both aging and obesity and decreased by exercise, suggesting that they could act as common mediators of EC dysfunction. We focused in this study on Serpinh1/Hsp47, as it is a collagen chaperone and has been shown to contribute to tissue fibrosis ^31, 45^, an important feature of many cardiac diseases. Recently, it was demonstrated in a mouse pressure overload model using Hsp47 cell type -specific knockout mice that Hsp47 in myofibroblasts is an important regulator of pathologic cardiac fibrosis ^45^. In line with our results in human cardiac ECs, collagen 1 production was decreased in the EC-specific Hsp47 deficient hearts ^45^. In human ECs, our results placed SERPINH1/HSP47 downstream of TGF-β and ROS, and demonstrated that its overexpression promoted mesenchymal features in human cardiac EC. Furthermore, SERPINH1/HSP47 was found to be important for extracellular collagen 1 deposition and EC proliferation/migration. Silencing of SERPINH1 also prevented the TGF-β induced appearance of TAGLIN-positive cells in human cardiac EC, which is considered as a marker for EndMT ^21, 37^. Based on the publicly available single-cell RNA sequencing data and immunohistochemistry of the human heart samples, SERPINH1/HSP47 is abundantly expressed in all cardiac endothelial populations. For further translational impact, the role of endothelial SERPINH1/HSP47 in aged, obese and hypertensive human hearts needs to be determined.

In conclusion, our data demonstrate that the major CVD risk factors significantly remodel the cardiac EC transcriptome promoting cell senescence, oxidative stress, TGF-β signaling and mesenchymal gene features, whereas exercise training provided opposite and protective effects. SerpinH1/Hsp47 was identified as one of the downstream effectors of TGF-β, which could provide a novel therapeutic target in endothelial cells.

## Materials and Methods

An expanded Methods section is available in the Supplementary material.

### Mouse Models

The committee appointed by the district of southern Finland approved all the animal experiments. Male C57BL/6J 7-8-week adult wild type mice were purchased from Janvier Labs, the detailed information about the mouse models, experimental procedures and treatments used in this study are described in the Data Supplement. The following four experimental groups were studied: exercise vs. sedentary, aged (18 mo) vs. young (2 mo), high-fat diet fed vs. chow-fed mice, and transverse aortic constriction (TAC) for two and seven weeks vs. sham-operated mice. Female C57BL/6J wild type mice of 19-24 months old were used for a separate exercise training experiment in old mice. The cohort size (n) for each experiment are shown in the respective figures or figure legends.

### Fluorescence-activated cell sorting of cardiac endothelial cells

The murine hearts were minced and incubated with 1mg/ml of collagenase type I, II and IV dissolved in DPBS containing 0.3mM CaCl2 at 37°C for 25 min. DMEM supplemented with 10% heat inactivated FCS was added and the cell suspension was filtered through 70μm nylon strainer. The cells were incubated with Fc receptor blocking antibody (CD16/32) for five minutes and followed by CD31, CD140a, CD45, Ter119 antibodies for 30 minutes. The live cardiac endothelial cells were defined as CD31^+^ CD45^-^ Ter119^-^ CD140a^-^ DAPI^-^ and the FACS Aria II (BD Biosciences) was used to gate, analyze and sort live cardiac endothelial cells. Data was acquired using FACS DIVA v8.0.1 and analyzed with FlowJo v10.1. The workflow is presented in detail in the **Figure S3A**.

### RNA seq data analysis of cardiac endothelial cells

FACS sorted cardiac ECs were homogenized using QIA shredder (Qiagen) and purified using RNeasy Plus Micro Kit (Qiagen). Prior to the library preparation step, RNA integrity and concentration of the samples were measured using Agilent Tape station and Qubit fluorescence assay, respectively. SMARTer Stranded Total RNA-Seq Kit V2 – Pico Input Mammalian (Takara Bio, USA) library preparation kit was used and 50M single end reads (1 x 75bp) were sequenced using illumina NextSeq 550 System. The sequenced reads were analyzed using Chipster high-throughput analysis software. The RNA sequencing data is deposited in GEO database, under the accession number GSE145263.

### Gene Ontology, Pathway analysis, Gene overlap and in silico gene characterization

To understand the biological role of differentially expressed genes, we used PANTHER classification system. VENNY 2.1 analysis software was used to identify overlapping genes and in silico gene characterization was performed using MetazSecKB knowledgebase, TargetP2.0 and SecretomeP1.0 softwares.

### Cell culture

Human cardiac arterial endothelial cells (HCAEC) or human umbilical vein endothelial cells (HUVEC) were transduced with lentiviral vectors (LV) encoding human SERPINH1-Myc overexpressing construct and four short hairpin gene silencing constructs for SERPINH1. The protocols for lentiviral vector production and gene transduction are explained in supplemental materials, and target sequences for shRNA constructs are listed in the **Supplementary Table 3**. For EndMT induction in HCAEC, a previously published method using TGF-β and H2O2 was used ^21, 37^.

### Immunohistochemistry, Real-Time Quantitative Polymerase chain reaction and Western Blotting

The detailed procedures for immunohistochemical stainings, western blotting and real-time qPCR are described in the Supplementary Materials and Methods section. Primer sequences for SYBR green and TaqMan real-time qPCR assays are listed in the **Supplementary Table 4**, and the antibodies used in immunohistochemistry and western blotting are listed in the **Supplementary Table 2**.

### Statistics

The data from the individual experiments were analyzed by student’s *t* test. P<0.05 value was considered statistically significant and P values in the graphs are mentioned as *P<0.05, **P<0.01 and ***P<0.001. The data is shown as mean ± SEM. The GraphPad Prism 7 software was used for statistical analysis. Statistics used for RNA sequencing data are described in detail in the Supplementary materials.

## Funding

We thank the Jenny and Antti Wihuri Foundation, Academy of Finland (RK, 297245), the Finnish Foundation for Cardiovascular Research (RK, KAH), the Sigrid Jusélius Foundation (RK), the Finnish Cultural Foundation (RK), The Finnish Medical Foundation (MIM), Biomedicum Helsinki Foundation (KAH) and Aarne Koskelo Foundation (KAH).

## Acknowledgements

We would like to thank Dr. Ralf Adams and Dr. Guillermo Luxán for their help in setting up the EC isolation. Dr. Seppo Kaijalainen for cloning the SERPINH1/HSP47 overexpression vector. Kirsi Mattinen, Päivi Leinikka, Maria Arrano de Kivikko, Ilse Paetau, Tanja Laakkonen and Tapio Tainola for their excellent technical help. We thank the Laboratory Animal Center, the Biomedicum Imaging Unit, the HiLife Flow Cytometry Unit, the Biomedicum Functional Genomics Unit, and the AAV Gene Transfer and Cell Therapy Core Facility for the help and facilities.

## Author contribution

KAH, EM and RK designed and KAH performed the experiments. KAH, SF, AA and RK analyzed the data. MIM collected the human heart samples. KAH and RK wrote the manuscript. All authors have seen, commented and accepted the manuscript.

## Competing interests

No competing interests.

---

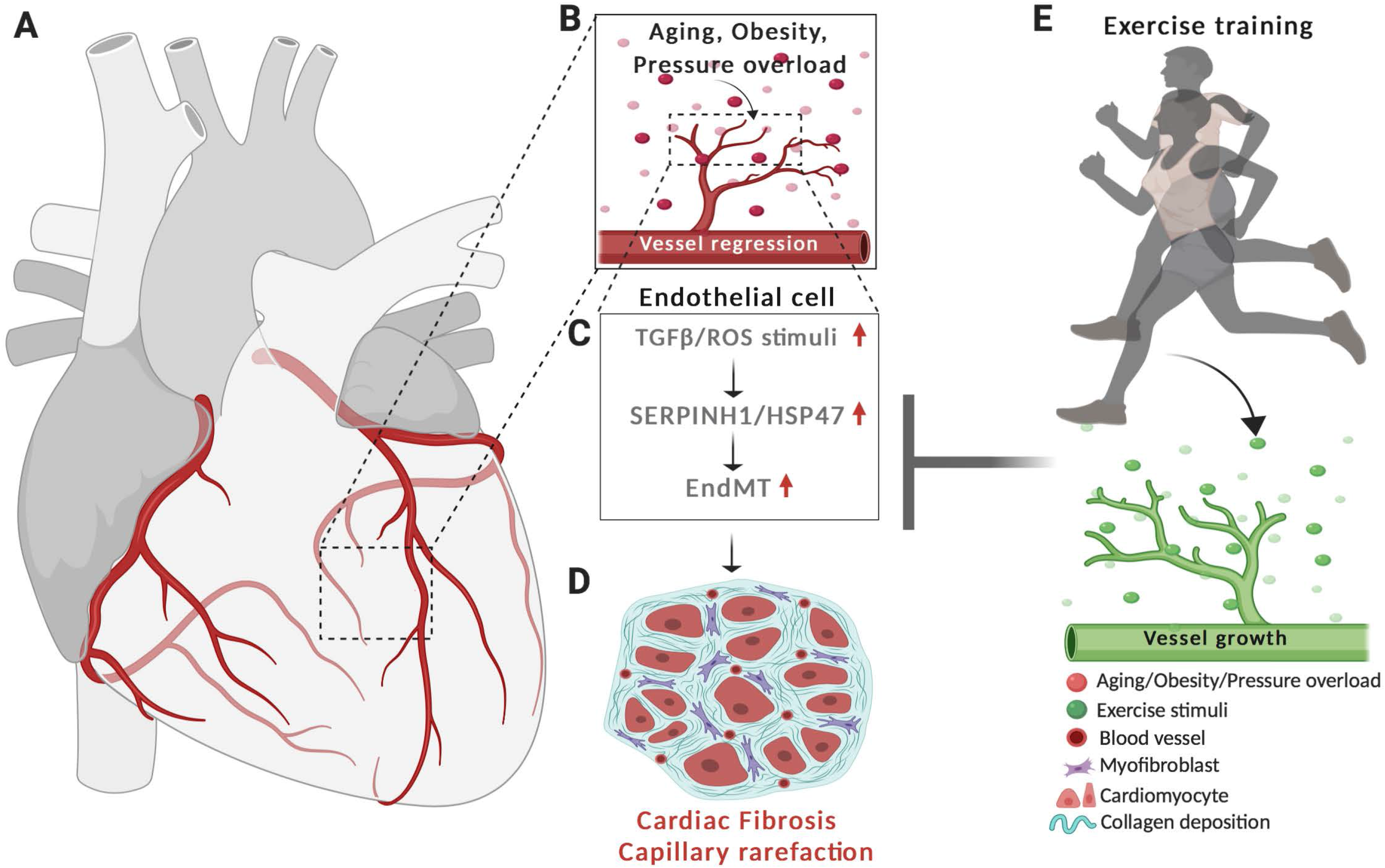
Graphical abstract demonstrating the cardiovascular disease risk factor mediated activation of TGF-β signaling and acquisition of mesenchymal features in cardiac EC. CVD risk factors aging, obesity and pressure overload trigger the regression of coronary vasculature by activating TGF-β/ROS signaling pathways and cellular senescence. These induce the expression of SerpinH1/Hsp47 and mesenchymal gene signature. SerpinH1/Hsp47 and EndMT are both involved in the development of tissue fibrosis by increasing collagen deposition in the extracellular matrix. Exercise training, in turn, increases coronary vasculature density, EC number and represses TGF-β signaling, mesenchymal gene expression and senescence related pathways.

## Supplementary Material

### Materials and Methods

#### Mouse Models

All animal experiments were approved by the committee appointed by the District of Southern Finland. Male C57BL/6J 7-8 -week adult wild type mice were purchased from Janvier Labs and used in the following experimental set-ups: physical activity (progressive exercise training vs sedentary), obesity (high-fat fed for 14 weeks vs chow), aging (18 months vs 2 months) and pressure overload/heart failure (transaortic constriction for two- and seven-weeks vs sham). Female C57BL/6J wild type mice of 19-24 months old were used for a separate exercise training experiment. The mice were housed in individually ventilated cages and acclimatized at least for one week in the animal facility before any experiments. The cohort size (n) for each experiment is indicated in the figures or figure legends.

#### Exercise Training

Ten-week-old C57BL/6J male mice or 19 to 24 months old female mice were trained on a treadmill (LE 8710, Bioseb). The mice were familiarized to the treadmill for three consecutive days with low speed (8-10 cm/s). Progressive training program consisted of 1-1.5 h training bouts five days a week for a total of six weeks with increasing speed, incline and/or duration each week. The following parameters in the treadmill controller were opted, tread inclination: 0°-10°; minimum and maximum tread speed: 10cm to 30cm per second; shock grid intensity: 0.2 mA. The aged mice were exercise-trained for four weeks and the same procedures were followed during the training program.

#### High Fat feeding

Ten-week-old C57BL/6J male mice were fed with standard chow diet or high-fat diet (HFD) containing 60% kcal derived from fat (Research Diets, D12492) for 4 or 14 weeks and used for immunohistochemistry or RNA-seq analysis, respectively.

#### Transverse Aortic Constriction Surgery

Ten-week-old C57BL/6J male mice were anesthetized with ketamine and xylazine. The mice were placed in supine position and intubated. The skin along the supra-sternal notch to mid sternum was incised to perform sternotomy to expose the aortic arch, right innominate and left common carotid arteries together with the trachea. Ligation of the transverse aorta between the right innominate left common carotid arteries against blunted 27-gauge needle with a 7-0 suture was performed and the needle was gently removed. The sternum and skin were ligated with monofilament polypropylene suture. Mice were placed in a warm chamber to recover, treated with analgesics (0.05mg/kg of Temgesic i.m.) at the time of the surgery and twice a day for following two days. For the control group (sham), all the steps in the surgical procedure were followed, except constricting the aorta. One group was killed two weeks and another group seven weeks after the surgery. Echocardiography was performed once a week during the experiment.

#### Echocardiography

To analyze cardiac function and ventricle dimensions, two-dimensional echocardiography images were acquired (Vevo 2100 Ultrasound, FUJIFILM Visual Sonics). The left ventricular internal diameter, left ventricular posterior wall thickness, interventricular septum thickness at end-systole and end-diastole were measured in M-mode along the parasternal short axis view, and analyzed by Simpson’s modified method ^1^.

#### Body Fat Measurement

The mice were anesthetized with ketamine and xylazine and the percentage of total body fat was measured using dual energy x-ray absorptiometry (Lunar PIXImus, GE Medical systems).

#### Oral Glucose Tolerance Test

Mice were fasted for four to five hours before the experiment. Glucose (1g/kg) was administered by oral gavage to mice. Blood from the tail tip was used to measure glucose levels at the following time points (15, 30, 60 and 90 min) using blood glucose meter (Contour, Bayer).

#### Immunofluorescent Staining

Frozen mouse heart sections (10μm) were cut with cryomicrotome and stained as described previously ^1^. The primary antibodies are listed in the **Supplementary Table 2**. Primary antibodies were detected with Alexa 488, 594 or 647 -conjugated secondary antibodies (Molecular Probes, Invitrogen). The sections were mounted with Vectashield Hard Set mounting media with DAPI (Vector Laboratories). The images were acquired with 20X, 40X air or 40x oil immersion objectives using AxioImager epifluorescent microscope (Carl Zeiss). The stained micrographs were initially adjusted for threshold, and an area fraction tool was used to quantify the area percentage of the vessels and collagen (Image J software, NIH).

#### Human Heart samples

Human heart samples were obtained from 4 organ donor hearts, which could not be used for transplantation e.g. due to size or tissue-type mismatch. The collection was approved by institutional ethics committee and The National Authority for Medicolegal Affairs.

#### Immunohistochemistry

The human paraffin heart sections (4μm) were cut, deparaffinized and rehydrated with xylene, descending concentration series of ethanol (99%, 95%, 70% and 50%) and H2O, and incubated in high pH antigen retrieval buffer containing 10 mM Tris, 1 mM EDTA, 0.05 % Tween 20 (pH 9.0). For HSP47 immunohistochemical analysis, VECTASTAIN Elite ABC kit (PK-6100) and DAB substrate was used to label and amplify the antibody signal. The 20X or 63X images were acquired with light microscope (Leica). For immunofluorescent staining, after the antigen retrieval step the sections were blocked with donkey immunomix (5% normal donkey serum, 0.2% BSA, 0.3% Triton X-100 in PBS), incubated overnight at 4°C with the primary antibodies for HSP47 and VE-Cadherin (CDH5) and detected with Alexa 488 and 594 conjugated secondary antibodies (Molecular probes, Invitrogen). The sections were mounted with Vectashield hardset with DAPI (Vector labs) and 40X images were acquired using AxioImager epifluorescent microscope (Carl Zeiss).

#### Isolation of Cardiac Endothelial Cells

The harvested hearts were briefly rinsed in ice-cold Dulbecco’s phosphate-buffered saline (DPBS, #14190-094, Gibco) supplemented with 0.3mM calcium chloride (CaCl2), cut opened longitudinally into two halves to expose the cardiac chambers and minced longitudinally and transversly into small pieces. To enzymatically dissociate the heart, 4ml of pre-warmed digestion media (1mg/ml of each collagenase types (type I (#17100-017), type II (#17101-015) and type IV (#17104-019) from Gibco were dissolved in DPBS containing 0.3mM CaCl2) and added to the minced hearts, incubated in water bath at 37°C for 25 min. During the digestion process, the samples were very gently mixed by vortexing for every 5 min. After incubation, the cell suspension was gently passed through T10 serological pipette 20 times. To neutralize the digestion, 10ml of rinsing media (Dulbecco’s modified eagle medium (#31053-028) supplemented with 10% heat inactivated FCS) was added to the cell suspension and filtered through the 70μm nylon cell strainer (Corning, #352350). Throughout the isolation process the cell suspensions were centrifuged for 5min, 300g and 4°C between each rinsing step. The cell pellet was resuspended in 5ml of ice-cold staining buffer (DPBS containing 2% heat inactivated FCS and 1mM EDTA). Before antibody staining, the cells were incubated with Fc receptor blocking antibody (CD16/32) for five minutes. The cells were incubated with the CD31, PDGFRa/CD140a, CD45, and Ter119 antibodies for 30 min (**Supplementary Table 2 for the antibody details)**. Prior to fluorescent associated cell sorting (FACS), the cells were rinsed twice with the staining buffer and filtered through 5ml cell strainer tubes (Corning, #352235).

#### Fluorescent Associated Cell Sorting (FACS)

The cells were passed through a 100μm nozzle. Multiple light scattering parameters for forward- and side-scatter properties of the cells were employed to gate, analyze and sort live cardiac endothelial cells. Initially, total cells were gated based on the forward and side-scatter area of the cells (FSC-A and SSC-A). The single cells were selected depending on forward scatter parameters area, height and width of the cells (FSC-A, FSC-H or FSC-W). DAPI was used to determine live and dead cells. To enrich and FACS sort pure and viable cardiac ECs, endothelial cells were stained with CD31, mesenchymal cells with PDGFRa/CD140a, leucocytes with CD45 and red blood cells with Ter119. The live cardiac endothelial cells were defined as CD31^+^ CD45^-^ Ter119^-^ CD140a^-^ DAPI^-^. Cells were sorted using FACS Aria II (BD Biosciences), the data was acquired with BD FACSDIVA v8.0.1 and further analyzed with FlowJo v10.1 (FlowJo, LLC) software. We verified the enrichment and purity of the FACS sorted Cardiac EC population (CD31+ PDGFRa (CD140a)-CD45-Ter119-DAPI-) by QPCR analysis for classical cardiac EC markers. Recently, we have used the same isolation method for single-cell RNAseq experiments, and these results show that there is about 3% contamination from other cells types, mainly pericytes and hemangioblasts.

#### RNA isolation

The sorted cardiac endothelial cells were immediately suspended in lysis buffer (350μl of RLT buffer plus 10μl of β-mercaptoethanol), the cells were homogenized in QIAshredder (#79654, Qiagen) and the RNA was purified using RNeasy Plus Micro Kit (#74034, Qiagen) according to the manufacturer’s instruction. The RNA integrity was analyzed with bioanalyzer (Agilent Tape Station 4200) and the concentration was determined by Qubit fluorescence assay (ThermoFisher). The cells from the post sort fractions were stained with propidium iodide (PI) and the viability of the cells were determined by Luna automated cell counter. The purity of the post sort fraction was determined by QPCR analysis for endothelial cell markers.

#### RNA sequencing of cardiac EC

Indexed cDNA library was synthesized using SMARTer Stranded Total RNA-Seq Kit V2 – Pico Input Mammalian (Takara Bio, USA) kit according to the manufacturer’s instructions. The library quality was determined using bioanalyzer, and sequenced using illumina NextSeq 550 System with the following specifications: 1 × 75bp, 50M single end reads were sequenced using NextSeq 500/550 High-Output v2.5 kit.

#### Differential gene expression

The sequenced reads were analyzed with the following software packages embedded in the Chipster analysis platform ^2^ (v3.12.2; https://chipster.csc.fi). Trimmomatic tool ^3^ (https://chipster.csc.fi/manual/trimmomatic.html) was used to preprocess Illumina single end reads. The HISAT2 package ^4^ (https://chipster.csc.fi/manual/hisat2.html) was employed to align the reads to mouse genome GRCm38.90 and the HTSeq count tool ^5^ (https://chipster.csc.fi/manual/htseq-count.html) to quantify the aligned reads per gene. The raw read count table for genes generated utilizing the HTSeq count were used as an input to perform two-dimensional principal component analysis (PCA) and unsupervised hierarchical clustering analysis using DESeq2 Bioconductor package ^6^ (https://chipster.csc.fi/manual/deseq2-pca-heatmap.html). Next, to perform the differential gene expression (DGE) analysis, the DESeq2 Bioconductor package ^6^ was used. The advantage of DEseq2 tool is sensitive and precise for analyzing the DEG in studies with few biological replicates. To reliably estimate the within group variance, Empirical Bayes shrinkage for dispersion estimation was used and a dispersion value for each gene was estimated through a model fit procedure (refer to the **Figure S5A**, which illustrates the shrinkage estimation for the experimental conditions). The gene features obtained after the dispersion estimation were used to perform statistical testing. Next, negative binomial generalized linear model was fitted for each gene and Wald test (raw p-value) was calculated to test the significance. Finally, DEseq2 applies Benjamini-Hochberg correction test to control the false discovery rate, FDR (refer to the **Figure S5B** indicating the distribution of raw and FDR adjusted pvalue for the experimental conditions). In our DEG analysis, we have set the FDR (p adj.) cut-off as less than or equal to 0.05 (FDR/p-adj ≤ 0.05) for pathway analysis and gene overlap analysis. The RNA sequencing data is deposited in the GEO database, under the series accession number **GSE145263**.

#### Gene Function and Pathway Analysis

The gene function and pathway analysis of the DGE were determined by performing statistical overrepresentation test using the PANTHER classification system ^7^ (V.14.1; http://www.pantherdb.org). The p<0.05 was considered for the further analysis and the data is presented as -log2(pvalue).

#### Gene Overlap and *in silico* Gene Characterization

The differentially expressed up- and downregulated genes (adjusted P-value 0.05) from the different experimental conditions were imported to VENNY 2.1 venn-diagram analysis software (BioinfoGP; https://bioinfogp.cnb.csic.es/tools/venny/) to identify genes which were significantly affected by several experimental conditions. The MetazSecKB knowledgebase^8^ (http://proteomics.ysu.edu/secretomes/animal/index.php), TargetP2.0 server ^9^ (http://www.cbs.dtu.dk/services/TargetP/index.php) and SecretomeP1.0 server ^10^ (http://www.cbs.dtu.dk/services/SecretomeP-1.0/) were used to characterize molecular functions, subcellular localizations and possible secretion properties of the identified common genes.

#### Cell Culture and Lentiviral Production

Human umbilical vein endothelial cells (HUVEC) and human cardiac arterial endothelial cells (HCAEC) were purchased from PromoCell. Both HUVEC and HCAEC were cultured and maintained in endothelial cell growth Basal Medium MV (C-22220, PromoCell) supplemented with Supplement Pack GM MV (C-39220, PromoCell) and gentamycin. For both gene overexpression and silencing studies, 80% confluent monolayer culture of HUVECs and HCAECs were used.

To overexpress SERPINH1 in EC, we cloned a lentiviral vector FUW-hSERPIH1-Myc (map and plasmid available by request). A scrambled sequence in the same vector was used as a control. 293FT cells (ATCC) were cultured and maintained in DMEM supplemented with 10% FCS and L-glutamine, and co-transfected with the lentiviral packaging plasmid vectors CMVg, CMVΔ8.9 and the target plasmid. The supernatants were collected at 48- and 72-hours, and concentrated by ultracentrifugation as described previously ^11^. For overexpression, HUVEC and HCAEC were transfected with lentivectors for 48 hours. For gene silencing studies, HCAEC were treated with lentivectors encoding for four independent clones of human shSERPINH1 for 24h. Subsequently, the cells were treated with puromycin (2ug/mL) for 48 hours to select the transduced cells. After selection, the cells were used for further analysis. The clone id and target sequence for human shSERPINH1 constructs are shown in the **Supplementary Table 3**.

#### Scratch wound assay

The SERPINH1 overexpressed or silenced HCAECs were seeded in the IncuCyte ImageLock 96-well microplate precoated with 0.1% gelatin and cultured in complete EC growth medium. To the confluent cell monolayers, 700 – 800 micron scratch wounds were introduced with IncuCyte WoundMaker, the wells were briefly rinsed with and maintained in complete EC growth medium. The kinetics of the cell migration were recorded and 10X phase contrast time-lapse images were acquired using IncuCyte Live-Cell Analysis System. The wound closure region was measured by Edge-detection and thresholding method in Image J software (NIH). The data is presented is as wound closure (%) relative to time.

#### EndMT assay

The coverslips or six well plates were precoated with 0.1% gelatin for 20min at 37°C, scrambled or SERPINH1 silenced HCAEC were seeded and cultured in complete EC growth medium. The cells were treated with or without 50ng/ml of recombinant human TGF-β (R&D Technologies) and/or 200μM hydrogen peroxide (Acros organics) for five days as described previously ^12, 13^.

#### Cell Staining

The cells grown on the coverslips were fixed with 4% PFA in PBS for 15 min. Blocking was done using donkey immunomix and the cells were stained with primary antibodies and secondary antibodies as indicated in the Supplemental Table 2. DAPI was used to stain the nucleus, and the cells were mounted using Vectashield (Vector labs). The amount of COL1 was quantified by adjusting 10X images for threshold and area fraction tool was used to quantify the area percentage of the collagen deposition (Image J software, NIH).

#### Real-Time Quantitative PCR

RNA from the cultured cells was purified and isolated using NucleoSpin RNA II Kit according to the manufacturer’s protocol (Macherey-Nagel). cDNA was synthesized with High-Capacity cDNA Reverse Transcription Kit (Applied biosystems, #4368814). SYBR green or TaqMan gene expression assays were performed using FastStart Universal SYBR green master mix (Sigma-Aldrich, #04913914001) and TaqMan gene expression master mix (Applied Biosystems, #4369016), respectively. mRNA expression was analyzed using Bio-Rad C1000 thermal cycler according to standardized protocol of the qPCR master mix supplier. The average of the technical triplicates for each sample was normalized to the housekeeping gene HPRT1. The mRNA expression levels were calculated and presented as fold change (Ctrl=1). The primer sequences are listed in the **Supplementary Table 4**.

#### Western Blotting

The cells were harvested and homogenized in lysis buffer containing 0.5%NP-40 (v/v) and 0.5%Triton X-100 (v/v) in PBS, supplemented with protease and phosphatase inhibitors (A32959, Pierce, Thermo Scientific). Protein concentration was determined using a BCA protein assay kit (Pierce, Thermo Scientific). Equal amounts of total protein were resolved in Mini-PROTEAN TGX Precast gels (Bio-Rad) and transferred to PVDF membrane (immobilon-P, Millipore). 5% BSA (wt/vol) and 0.1% Tween 20 (v/v) in TBS was used to block the membranes followed by incubation with primary antibodies (**Supplementary Table 2**) overnight at 4°C. HRP-conjugated secondary antibodies (DAKO) were used, and HRP signals were developed with Super-Signal West Pico Chemiluminescent substrate or Femto Maximum sensitivity substrate (Thermo Scientific). The blots were imaged with Odyssey imager (Li-COR Biosciences) or Chemi Doc imaging system (Bio-Rad) and quantified with Image Studio Lite Software (Li-COR Biosciences).

#### Statistics

The data from the individual experiments were analyzed by student’s *t* test. P<0.05 value was considered statistically significant and P values in the graphs are shown as *P<0.05, **P<0.01 and ***P<0.001. The data is shown as mean ± SEM. The GraphPad Prism 7 software was used for statistical analysis.

---

**Figure S1.**
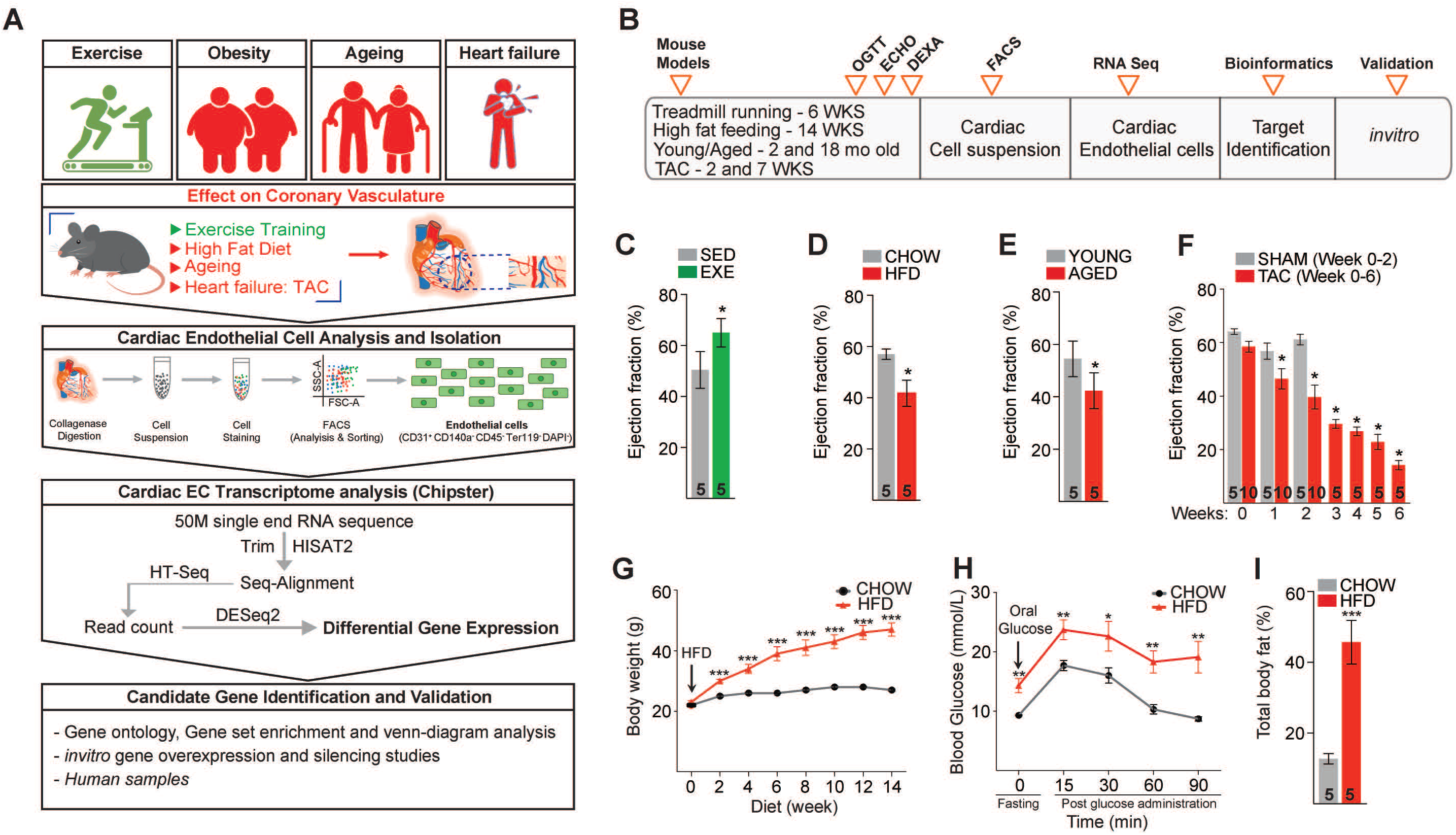
Schematic of the experimental set-up to elucidate the impact of cardiovascular disease risk factors on cardiac endothelial cell transcriptome and the validation of the experimental CVD risk factor models. **A.** Experimental workflow demonstrating the mouse models used to mimic CVD risk factors in C57BI/6J mice, analysis and isolation of cardiac ECs by fluorescence-activated cell sorting, bioinformatic analyses of the cardiac EC transcriptome, identification and validation of candidate genes using human ECs and heart tissue. **B.** Experimental timeline of exercise training (6 weeks of treadmill running), high fat diet (14 weeks of high fat feeding), physiological ageing (18 months old) and pressure overload -induced heart failure by transaortic constriction in mice. **C-F.** Ejection fraction in each of the four experiments. **G.** Body weight (g) during the HFD experiment. **H.** Blood glucose levels during oral glucose tolerance test (mmol/L), and I. total body fat (%) measured after 14 weeks of high-fat diet. Data is presented as mean ± SEM. Student’s t test was used, *p<O.05, **p<O.01, ***p<O.001 (In panel **C-F and I**, number of mice in each experimental group are indicated in the respective graph, In panel **G-H**, N=4-5 mice/group were analysed.

**Figure S2.**
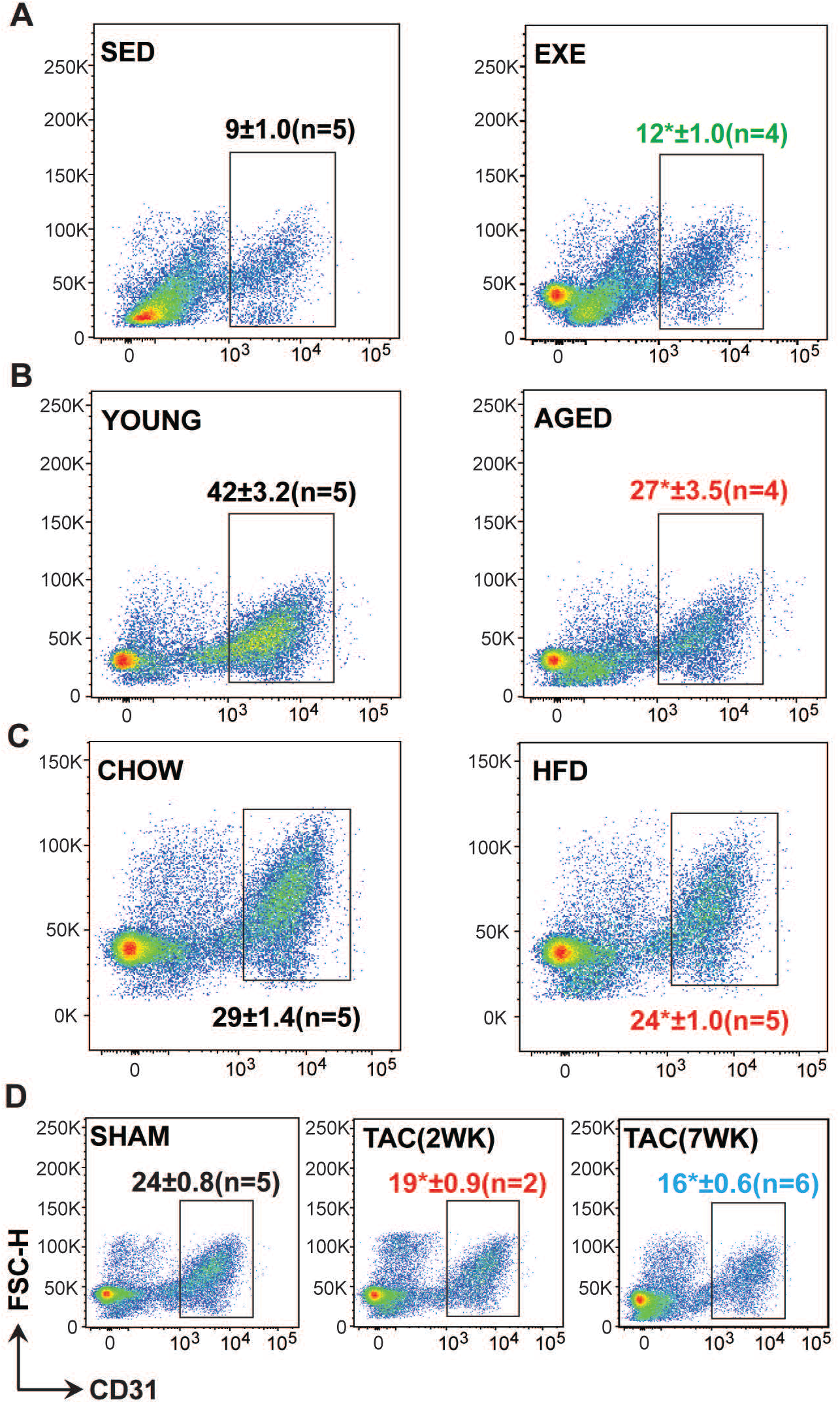
FACS analysis of cardiac EC. **A-D.** Representative pseudocolor FACS plots showing the gating and percentage of cardiac ECs (CD31+ CD140a-CD45-Ter119-DAPI-) in the different treatment groups. In panel **A-D,** number of mice in each experimental group are indicated in the respective FACS plots.

**Figure S3.**
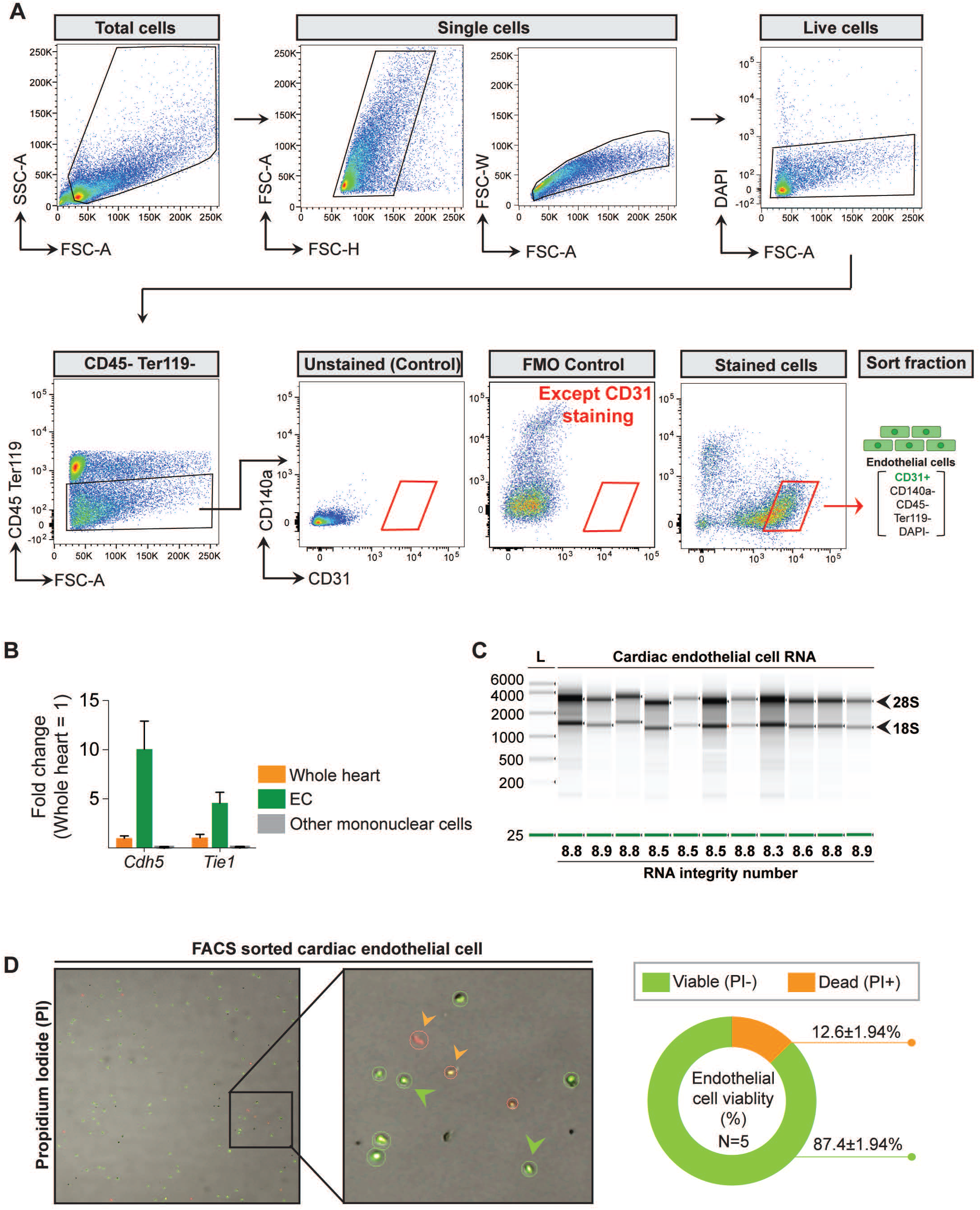
Quality metrics of the FAC5 sorted cardiac EC. **A.** Gating strategy to sort cardiac ECs (CD31+ CD140a-CD45-Ter119-DAPI-). **B.** Purity analysis of the post sort EC fraction by QPCR (N=4 mice/group were analyzed). **C.** Representative image of the bioanalyzer data showing the RIN values of the isolated RNA. **D.** Mononuclear cells and pie chart of the post sort EC fraction showing viable cells in green and dead cells in red.

**Figure S4.**
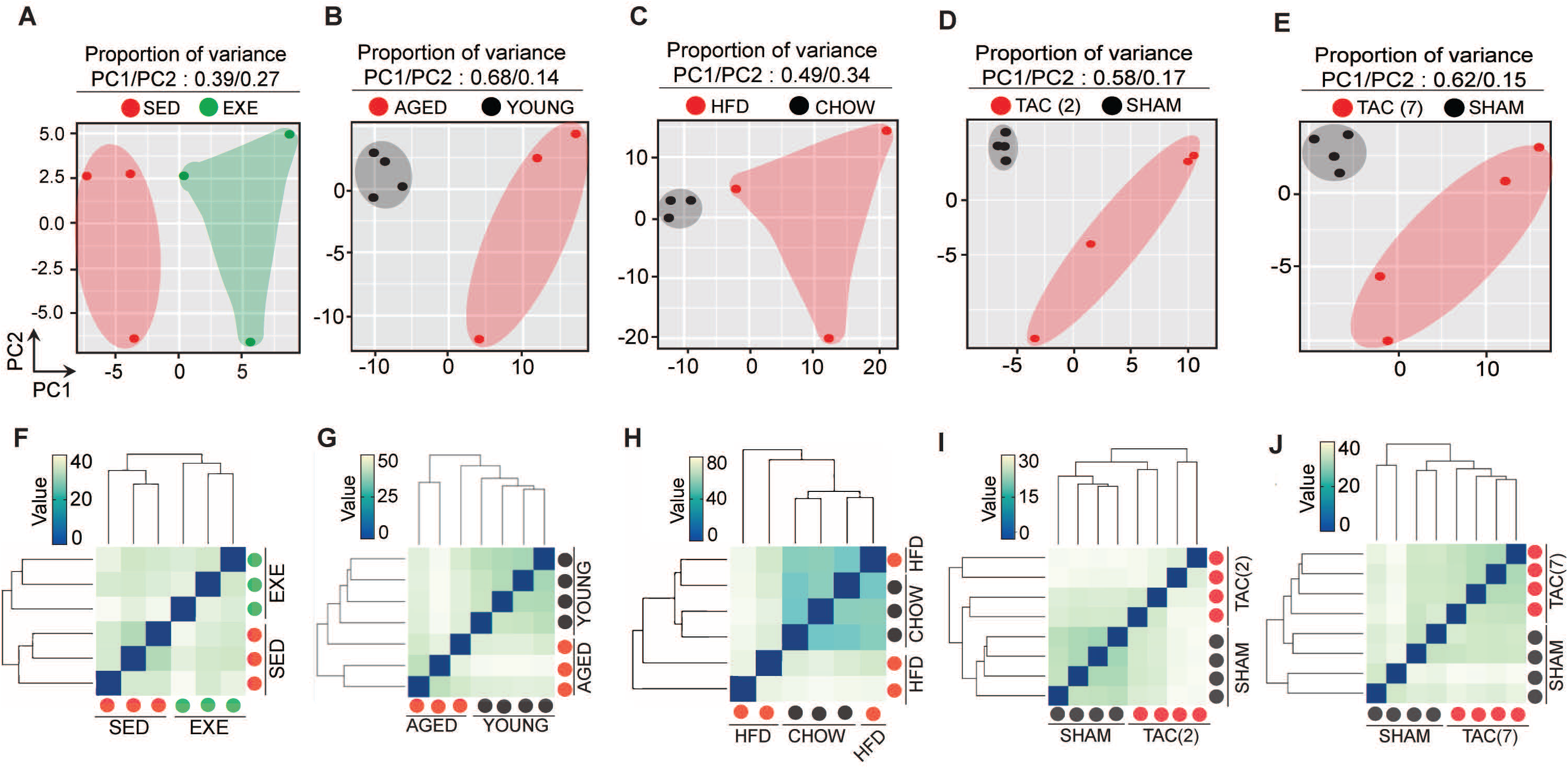
PCA plot and unsupervised hierarchical clustering of cardiac endothelial cell transcriptome from exercise trained, aged, obese and TAC-treated mice. **A-E.** Two dimensional principal component analysis, **F-J.** Unsupervised hierarchical clustering of cardiac endothelial transcriptome in the indicated experimental group. Eachcolor coded circles (Red, Green and Black) in the PCA and Unsupervised clustering plot indicate one biological sample and N=3-4 mice/experimental condition were analysed.

**Figure S5.**
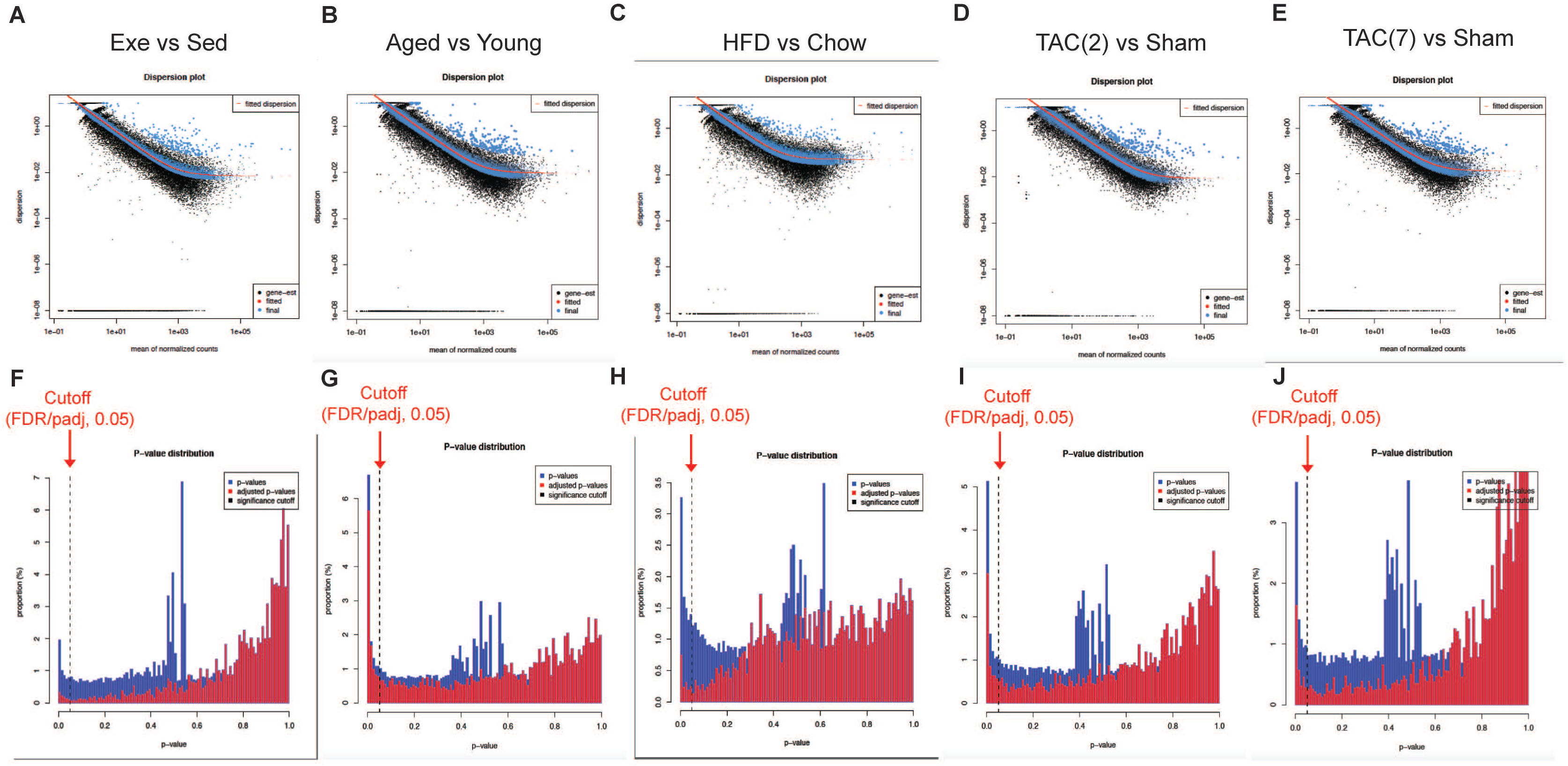
Dispersion mean plot and p-value distribution plot of the indicated RNA sequencing experiments. **A-E.** Plot of dispersion estimates at different count levels, showing black dot (Dispersion estimate for each gene as obtained by considering the information from each gene separately), Red line (Fitted estimates showing the dispersions’ dependence on the mean), Blue dot (The final dispersion estimates shrunk from the gene-wise estimates towards the fitted estimates. The values are used for further statistical testing). Blue circles (Genes which have high gene-wise dispersion estimates and are hence labelled dispersion outliers and not shrunk toward the fitted trend line. **B.** Plot of the raw p-value (Wald test) indicated in blue bar and the false discovery rate distribution or adjusted p-value (Benjamini-Hochberg adjusted p-value) of the statistical test. The arrow indicates cutt-off point False discovery rate thereshold of 0.05 and the genes with FOR values less than or equal to the cutoff points were used for futher analysis.

**Figure S6.**
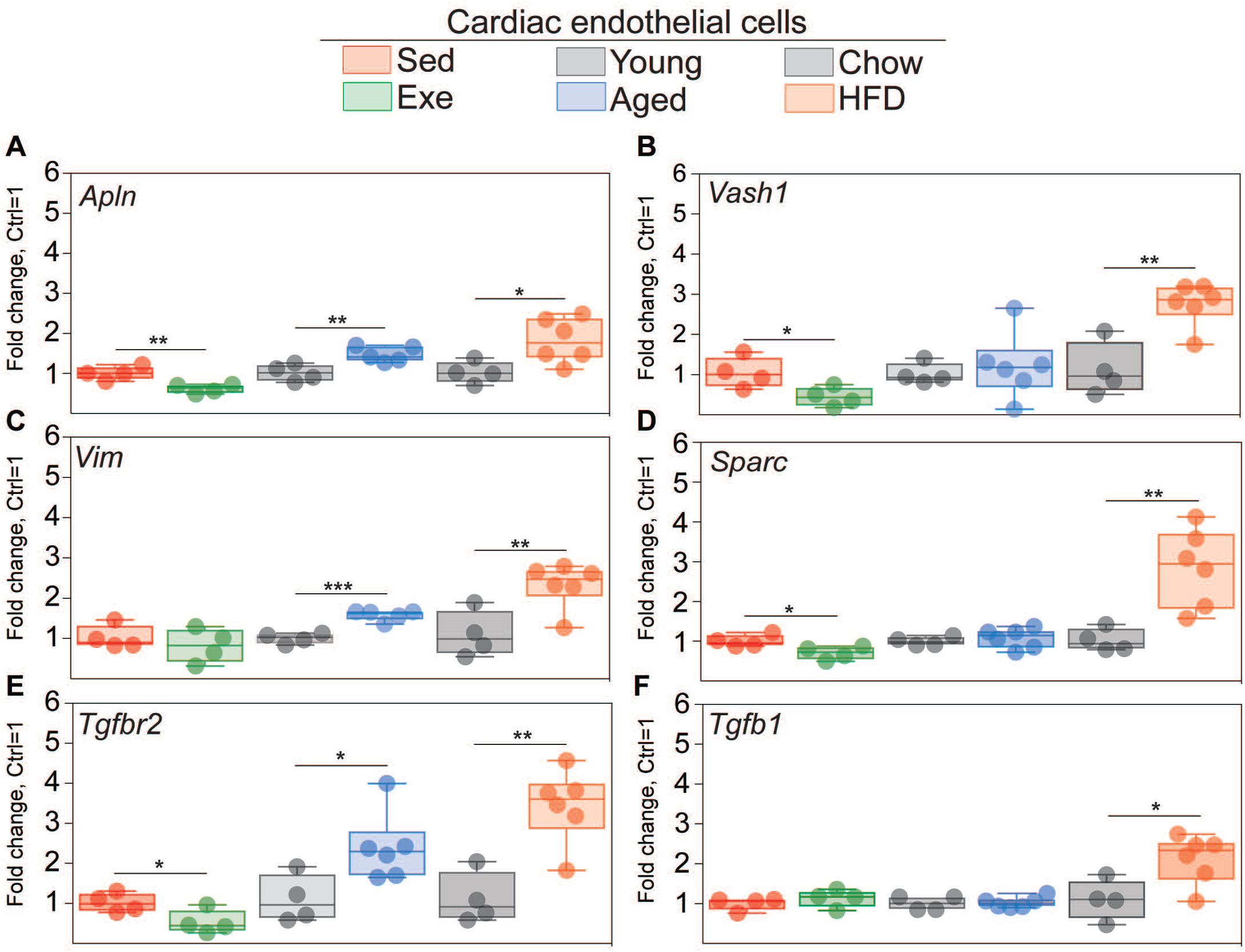
QPCR validation of the indicated genes in the cardiac EC during aging, obesity, exercise training in young and aged mice. A-F. mRNA expression of Apln, Vim, Tgfbr2, Vash1, Sparc and Tgfb1 in the cardiac EC of indicated experimental groups (N=4-6/group). Gene expression is normalised to HPRT1 expression. Data is presented as mean ± SEM. Student’s t test was used, · p<O.05, ··p<O.01, ···p<O.001.

**Figure S7.**
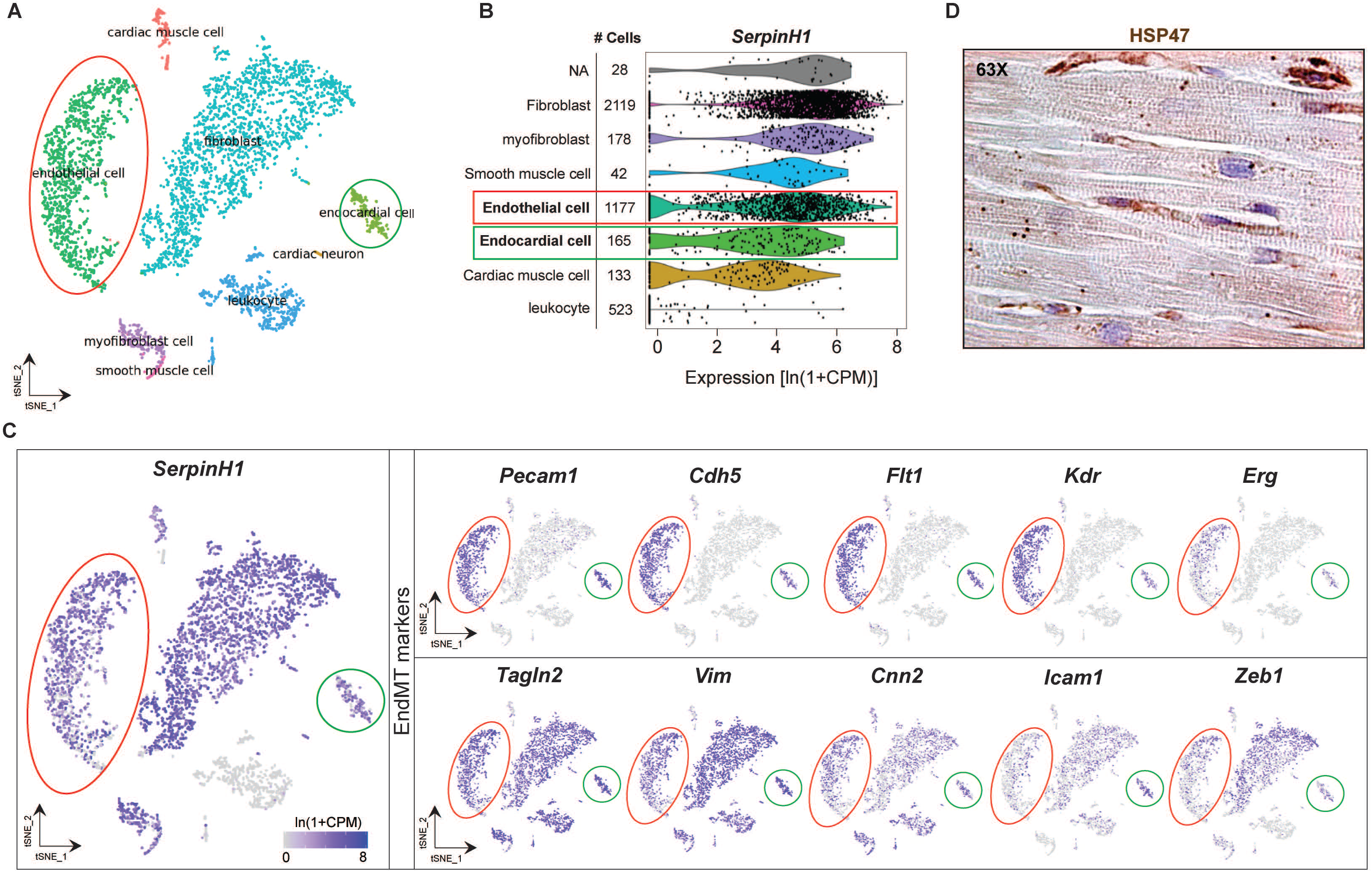
Expression of SerpinH1/HSP47 in different cardiac cell types and in the human heart. A tSNE plot showing **A.** cardiac cell types in the adult mouse heart. **B.** Violin plots showing the levels of SerpinH1 transcripts in fibroblasts, myofibroblasts, smooth muscle cells, endothelial cells, endocardial cells, cardiac muscle cells and leucocytes (each black dot denotes a single cell). **C.** tSNE plot showing the expression of SerpinH1 and EndMT genes in the endothelial cell cluster (cells within red circle) and endocardial cluster (cells within green clusters) Illustrations in the panels A-E were analyzed and acquired from publicly available,:5,i!igle cell database Tabula muris: https://tabula-muris.ds.czbiohub.org/ . **D.** Representative longitudinal IHC image of human heart demonstrating strong SERPINH1/HSP47 expression in interstitial cells (fibroblasts, endotheliaR!ells) and weak staining within cardiomyocytes. Scale bar 100 µm.

**Figure S8.**
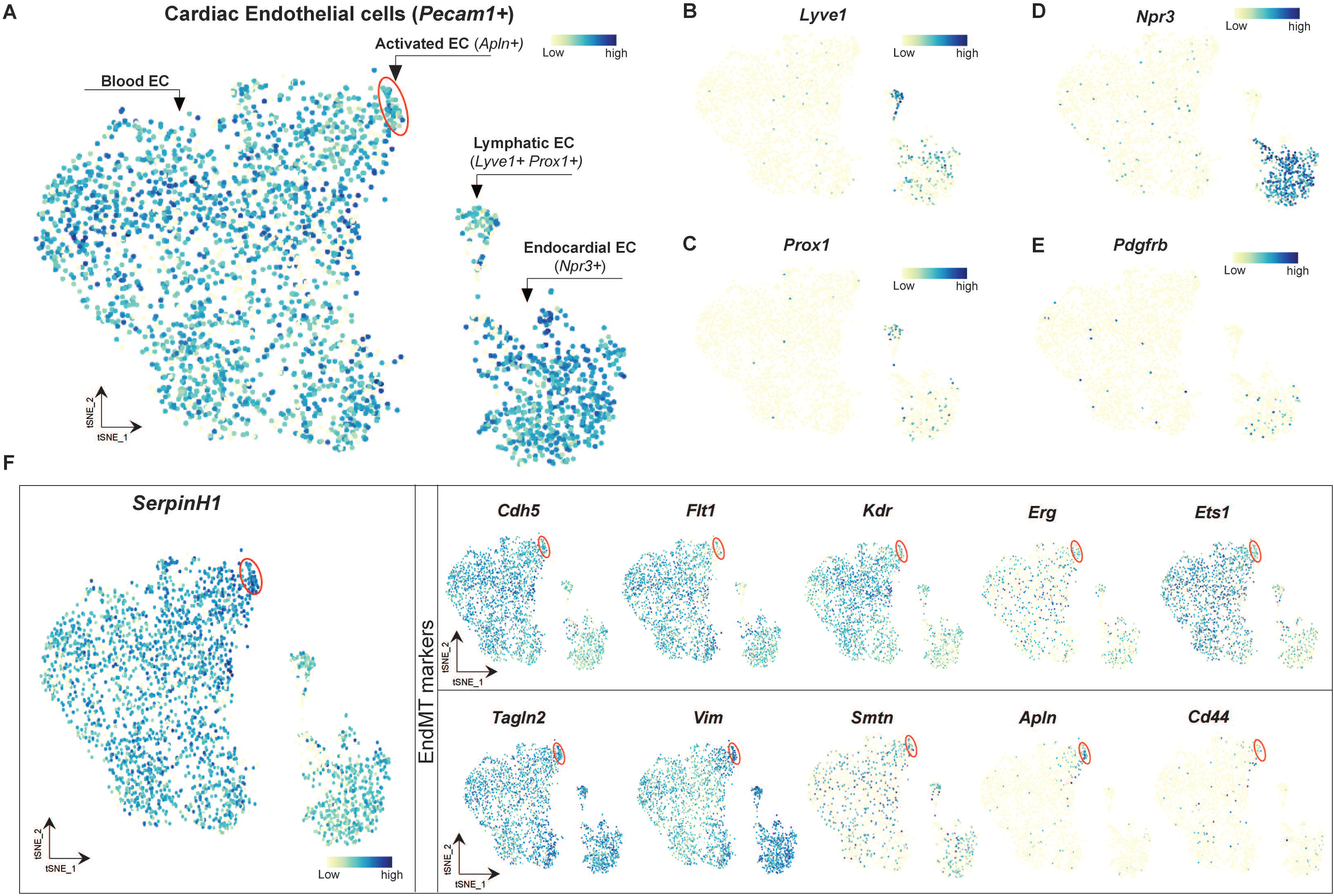
SERPINH1 expression in different subset of cardiac EC. **A-D.** t-SNE plot showing the Blood EC, Activated EC (Apln+) highlighted within red circle, lymphatic EC (lyve1+ Prox1+) and endocardial EC (Npr3+), **E.** Pdgfrb expr on in cardiac EC population, **F.** SerpinH1 is expressed in all endothial cell population, highly expressed in Apln+ activated EC (cell cluster within red circle) which shows decreased expression of EC genes (Cdh5, Flt1, Kdr, Erg and Ets1) and ina’e’ased expression of Mesenchymal genes (Tagln2, Vim, Smtn, Cd44). Illustration in the panel A-F were analysed from publicly available heart EC atlas from peter carmeliet lab (https://endotheliomics.shinyapps.io/ec_atlas/).

**Figure S9.**
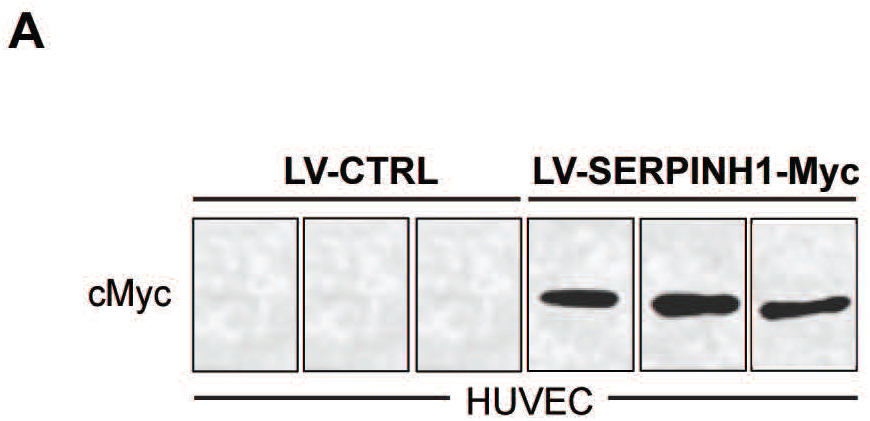
5ERPINH1 overexpression in endothelial cells. **A.** Western blot analysis of cMyc expression in LV-SERPINH1-Myc treated HUVEC (N=3 biological replicates/group).

**Supplementary Table 1.** Echocardiography measurements of cardiac function and ventricular dimensions.

| Experiment groups | IVS,d<br>(mm) | IVS,s<br>(mm) | LVID,d<br>(mm) | LVID,s<br>(mm) | LVPW,d<br>(mm) | LVPW,s<br>(mm) | LV Vol,d<br>(μl) | LV Vol,s<br>(μl) | LV mass<br>(mg) | FS<br>(%) |
| --- | --- | --- | --- | --- | --- | --- | --- | --- | --- | --- |
| **6 weeks of treadmill running** |  |  |  |  |  |  |  |  |  |  |
| EXE (n=5); 3 mo old | 1.02±0.14 | 1.33±0.13 | 3.84±0.10 | 2.56±0.12<sup>†</sup> | 0.74±0.05 | 1.25±0.05 | 63.8±4.01 | 23.9±2.73<sup>†</sup> | 103.6±19.1 | 33.5±2.3 |
| SED (n=5); 3 mo old | 0.89±0.07 | 1.13±0.11 | 3.93±0.05 | 2.94±0.10 | 0.89±0.09 | 1.22±0.13 | 67.2±1.85 | 33.5±2.83 | 106.9±4.4 | 25.3±1.9 |
| **14 weeks of high fat diet feeding** |  |  |  |  |  |  |  |  |  |  |
| HFD (n=5); 3 mo old | 1.11±0.04<sup>†</sup> | 1.39±0.06 | 3.92±0.13 | 3.10±0.21 | 0.86±0.03<sup>†</sup> | 1.09±0.08 | 67.2±5.32 | 39.3±6.5 | 121.7±5.2 | 21.1±3.3<sup>†</sup> |
| CHOW (n=5); 3 mo old | 0.93±0.05 | 1.41±0.03 | 4.19±0.08 | 2.96±0.08 | 0.75±0.04 | 1.05±0.15 | 78.5±3.54 | 34.2±2.4 | 107.9±3.8 | 29.4±1.6 |
| **18 months of aging** |  |  |  |  |  |  |  |  |  |  |
| AGED (n=6); 18 mo old | 1.05±0.03<sup>†</sup> | 1.39±0.06<sup>†</sup> | 4.16±0.11 | 3.31±0.13<sup>†</sup> | 1.09±0.06<sup>†</sup> | 1.34±0.06<sup>†</sup> | 77.2±4.4 | 44.9±4.1<sup>†</sup> | 150.1±9.1<sup>†</sup> | 20.7±1.6<sup>†</sup> |
| YOUNG (n=6); 2 mo old | 0.85±0.04 | 1.19±0.06 | 3.94±0.10 | 2.84±0.13 | 0.77±0.06 | 1.13±0.03 | 68.1±4.4 | 31.0±3.3 | 93.8±3.9 | 27.9±1.7 |
| **Trans-aortic constriction: Week 0-6** |  |  |  |  |  |  |  |  |  |  |
| Sham(n=5);Week 0 | 0.78±0.03 | 1.12±0.03 | 3.50±0.06 | 2.31±0.05 | 0.79±0.01 | 1.19±0.03 | 51.0±2.0 | 18.2±1.2 | 73.5±1.6 | 34.1±0.8 |
| TAC (n=10);Week 0 | 0.77±0.03 | 1.10±0.04 | 3.67±0.12 | 2.57±0.13 | 0.81±0.02 | 1.20±0.03 | 58.1±5.0 | 24.8±3.6 | 80.1±3.1 | 30.4±1.3 |
| Sham(n=5);Week 1 | 0.73±0.03 | 1.07±0.06 | 3.88±0.09 | 2.74±0.12 | 0.75±0.03 | 1.09±0.05 | 65.4±3.7 | 28.4±3.1 | 80.1±2.2 | 29.4±1.9 |
| TAC (n=10);Week 1 | 0.87±0.04<sup>†</sup> | 1.18±0.06 | 3.92±0.12 | 3.03±0.18 | 0.98±0.06<sup>†</sup> | 1.25±0.07 | 67.6±5.1 | 37.8±5.0 | 111.6±7.1<sup>†</sup> | 23.2±2.3 |
| Sham(n=5);Week 2 | 0.70±0.04 | 1.07±0.05 | 3.89±0.06 | 2.63±0.07 | 0.76±0.02 | 1.11±0.04 | 65.6±2.8 | 25.6±1.8 | 79.3±3.2 | 32.4±1.4 |
| TAC (n=10);Week 2 | 0.87±0.04<sup>†</sup> | 1.14±0.05 | 4.12±0.15 | 3.34±0.21<sup>†</sup> | 1.02±0.03<sup>†</sup> | 1.27±0.04<sup>†</sup> | 76.9±7.3 | 48.0±7.4<sup>†</sup> | 124.4±6.2<sup>†</sup> | 19.5±2.6<sup>†</sup> |
| TAC (n=5); Week 3 | 0.97±0.04<sup>†</sup> | 1.17±0.04 | 4.38±0.19<sup>†</sup> | 3.77±0.16 | 1.08±0.06<sup>†</sup> | 1.29±0.07 | 87.8±9.8 | 61.4±6.6<sup>†</sup> | 153.6±12.2<sup>†</sup> | 13.8±0.9<sup>†</sup> |
| TAC (n=5); Week 4 | 0.92±0.05<sup>†</sup> | 1.20±0.06 | 4.60±0.23<sup>†</sup> | 4.04±0.24 | 1.25±0.10<sup>†</sup> | 1.37±0.09 | 99.0±13<sup>†</sup> | 72.8±11<sup>†</sup> | 178.3±9.9 | 12.4±0.8<sup>†</sup> |
| TAC (n=5); Week 5 | 0.89±0.10<sup>†</sup> | 1.15±0.11 | 4.92±0.07<sup>†</sup> | 4.40±0.09 | 1.00±0.05<sup>†</sup> | 1.12±0.03<sup>†</sup> | 114.0±4<sup>†</sup> | 87.8±4.3<sup>†</sup> | 165.6±10 | 10.6±1.4<sup>†</sup> |
| TAC (n=5); Week 6 | 0.81±0.06 | 0.96±0.08 | 5.15±0.22<sup>†</sup> | 4.82±0.24 | 0.98±0.09<sup>†</sup> | 1.01±0.09 | 127.8±13<sup>†</sup> | 110.2±13<sup>†</sup> | 164.9±7.9 | 6.4±0.8<sup>†</sup> |

mo: Months; **IVS,d**: Interventricular septum thickness at end-diastole; **IVS,s**: Interventricular septum thickness at end-systole; **LVID,d**: Left ventricular internal dimension at end-diastole; **LVID,s**: Left ventricular internal dimension at end-systole; **LVPW,d**: Left ventricular posterior wall thickness at end-diastole; **LVPW,s**: Left ventricular posterior wall thickness at end-systole; **LV Vol, d**: Left ventricular volume at end-diastole; **LV Vol,s**: Left ventricular volume at end-systole; **LV mass**: Left ventricular mass; **FS**: Fractional shortening.
Data are mean ± SEM. Student *t* test was used, <sup>†</sup> *p*<0.05.

**Supplementary Table 2:** List of antibodies used for fluorescence-activated cell sorting (FACS), immunofluorescent staining (IF) and Western blotting (WB).

| Antibody | Host | Application | Company/Catalog number |
| --- | --- | --- | --- |
| FITC-CD31 | Mouse | FACS | Invitrogen/RM5201 |
| Pacificblue-CD45 | Mouse | FACS | Biolegend/103125 |
| Pacificblue-Ter119 | Mouse | FACS | Biolegend/116231 |
| PE-Cyanine7-CD140a | Mouse | FACS | eBioscience/25-1401 |
| CD16/CD32 (Fc blocker) | Mouse | FACS | BD Biosciences/553142 |
| Mouse CD31 | Rat | IF | BD Pharmingen/553370 |
| Human VEcadherin | Rabbit | IF | Cell Signaling/#2500S |
| Tagln | Sheep | WB/IF | R&D Biosystems/AF7886 |
| c-MYC | Mouse | WB/IF | Thermo Fisher/13-2500 |
| HSP47 | Mouse | WB/IHC/IF | Enzo Life Sciences/ADI-SPA-470-D |
| Collagen 1 | Rabbit | IF | Abcam/ab34710 |
| aSMA | Mouse | IF | Sigma-Aldrich/A5228 |
| GAPDH | Mouse | WB | Sigma-Aldrich/CB1001 |

**Supplementary Table 3:**
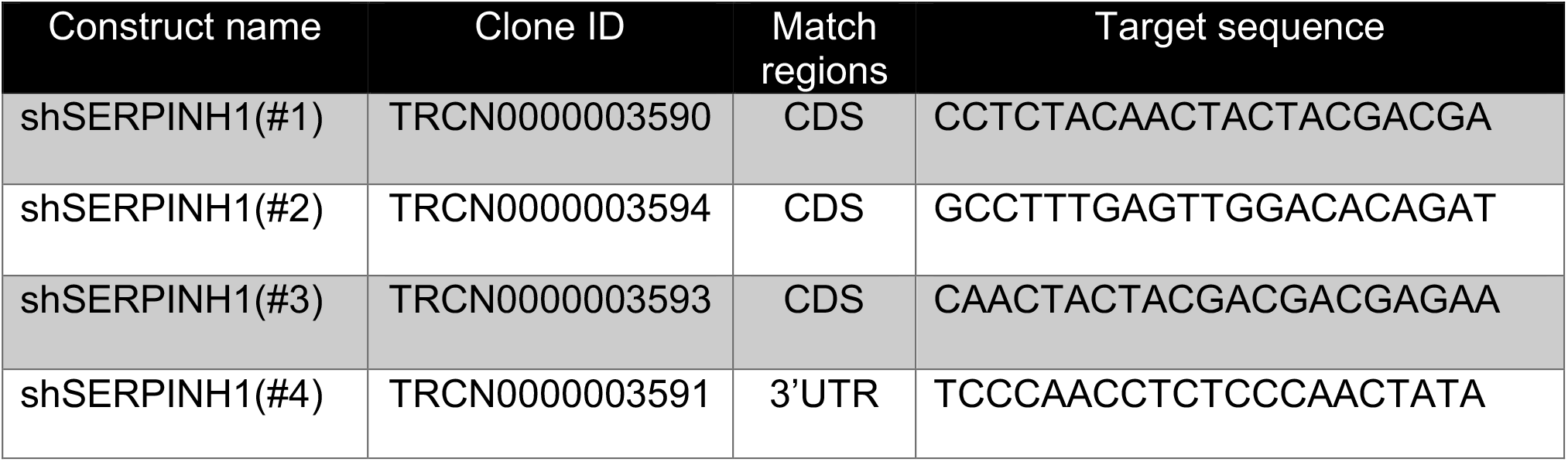
List of human shRNA constructs used in the gene silencing studies.

**Supplementary Table 4:**
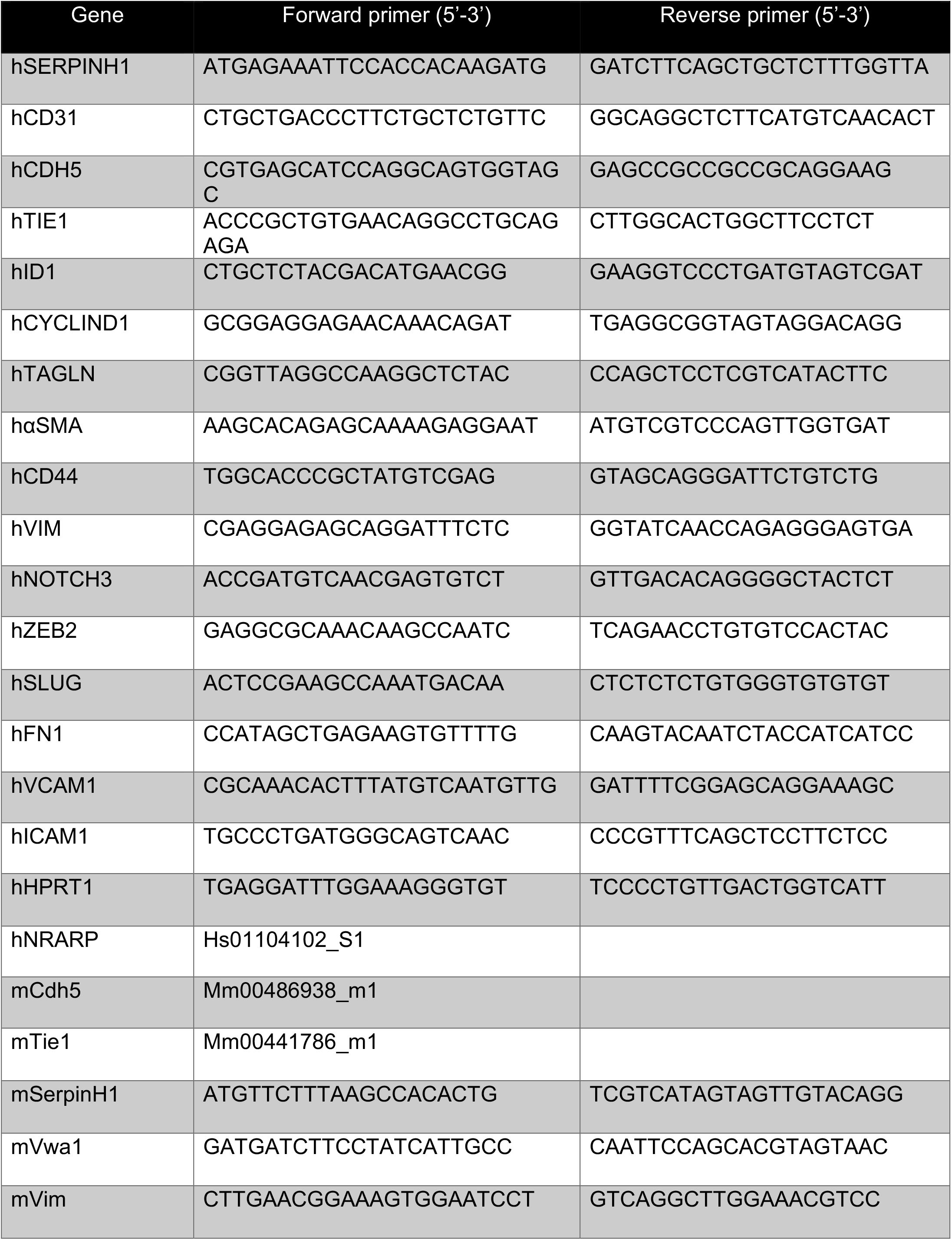
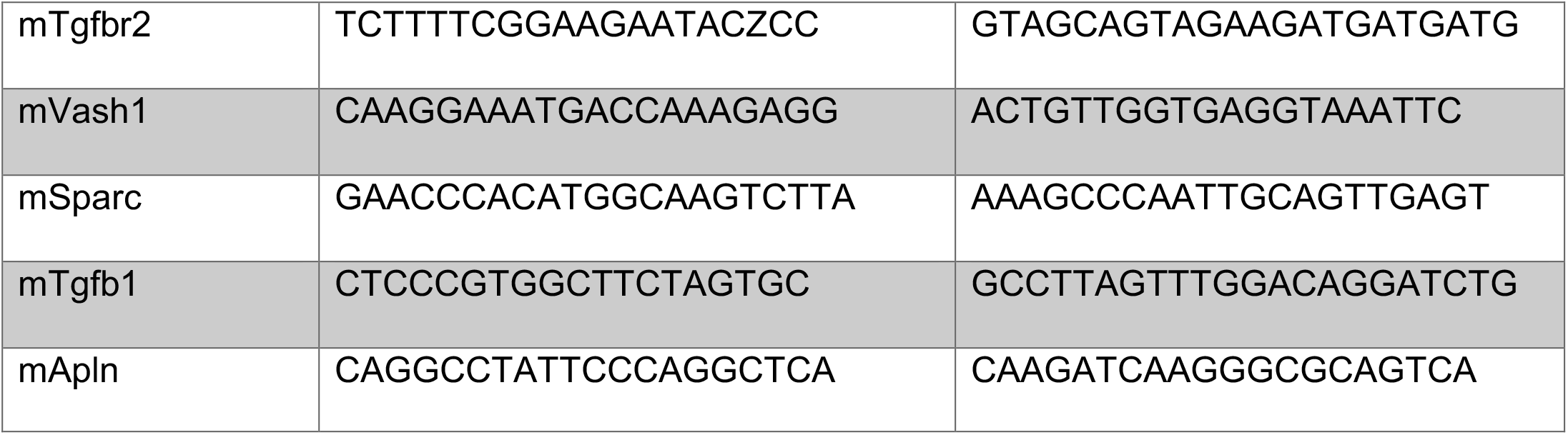
Primer pair sequences for SYBR green and TaqMan real-time qPCR assays.

**Supplementary Table 5A:** Reference list for endothelial and mesenchymal genes indicated in the Figure 4B (EXE vs. SED) heat map.

| Gene | Description | Reference # |
| --- | --- | --- |
| *Malat1* | Metastasis Associated Lung Adenocarcinoma Transcript 1 | 14 |
| *Mgp* | Matrix Gla Protein | 15 |
| *Ankrd11* | Ankyrin Repeat Domain 11 | 16 |
| *Plcb4* | Phospholipase C Beta 4 | 17 |
| *Rock1* | Rho Associated Coiled-Coil Containing Protein Kinase 1 | 18 |
| *Adgrg6* | Adhesion G Protein-Coupled Receptor G6 | 19 |
| *Ppp4r3a* | Protein Phosphatase 4 Regulatory Subunit 3A | 20 |
| *Phip* | Pleckstrin Homology Domain Interacting Protein | 21 |
| *Clk1* | CDC Like Kinase 1 | 22 |
| *Ankrd12* | Ankyrin Repeat Domain 12 | 23 |
| *Ppl* | Periplakin | 24 |
| *Krit1* | KRIT1 Ankyrin Repeat Containing | 25, 26 |
| *Calcrl* | Calcitonin Receptor Like Receptor | 27 |
| *Apln* | Apelin | 28 |
| *Aplnr* | Apelin Receptor | 29 |
| *Vash1* | Vasohibin 1 | 30, 31 |
| *Fscn1* | Fascin Actin-Bundling Protein 1 | 32 |
| *Cd93* | CD93 Molecule | 33, 34 |
| *Vwa1* | Von Willebrand Factor A Domain Containing 1 | 35, 36 |
| *Adamts4* | ADAM Metallopeptidase With Thrombospondin Type 1 Motif 4 | 37 |
| *Sparc* | Secreted Protein Acidic And Cysteine Rich | 38 |
| *Col4a2* | Collagen Type IV Alpha 2 Chain | 39, 40 |
| *Cd34* | CD34 Molecule | 41, 42 |
| *Tuba1a* | Tubulin Alpha 1a | 43 |
| *Cx3cl1* | C-X3-C Motif Chemokine Ligand 1 | 44 |
| *Mest* | Mesoderm Specific Transcript | 45 |
| *Cnn2* | Calponin 2 | 46 |
| *Cd44* | CD44 Molecule (Indian Blood Group) | 47 |
| *Trp53* | Tumor Protein P53 | 48 |
| *Prex1* | Phosphatidylinositol-3,4,5-Trisphosphate Dependent Rac Exchange Factor 1 | 49 |
| *Hspg2* | Heparan Sulfate Proteoglycan 2 | 29 |
| *Tnfaip1* | TNF Alpha Induced Protein 1 | 50 |
| *Lamb1* | Laminin Subunit Beta 1 | 51 |
| *Ltbp4* | Latent Transforming Growth Factor Beta Binding Protein 4 | 52 |
| *Sept4* | Septin 4 | 53 |
| *Unc5b* | Unc-5 Netrin Receptor B | 54 |

**Supplementary Table 5B:** Reference list for endothelial and mesenchymal genes indicated in the Figure 4C (Aged vs. Young) heat map.

| Gene | Description | Reference # |
| --- | --- | --- |
| *Npr3* | Natriuretic Peptide Receptor 3 | 55, 56 |
| *Tagln2* | Transgelin 2 | 57 |
| *Vim* | Vimentin | 58 |
| *Msn* | Moesin | 59-61 |
| *Socs5* | Suppressor Of Cytokine Signaling 5 | 62 |
| *Vcam1* | Vascular adhesion molecule 1 | 63 |
| *Notch4* | Notch Receptor 4 | 41, 64 |
| *Stat6* | Signal Transducer And Activator Of Transcription 6 | 65 |
| *Adamtsl4* | ADAMTS Like 4 | 66 |
| *Jag2* | Jagged Canonical Notch Ligand 2 | 41 |
| *Cdh13* | Cadherin 13 | 67, 68 |
| *Hif1a* | Hypoxia Inducible Factor 1 Subunit Alpha | 41 |
| *Ism1* | Isthmin 1 | 69, 70 |
| *Tbx3* | T-Box Transcription Factor 3 | 71 |
| *Tgfa* | Transforming Growth Factor Alpha | 72 |
| *Furin* | Furin, Paired Basic Amino Acid Cleaving Enzyme | 73 |
| *Eng* | Endoglin | 74, 75 |
| *Smad6* | SMAD Family Member 6 | 76, 77 |
| *Smad7* | SMAD Family Member 7 | 76, 77 |
| *Sox17* | SRY-Box Transcription Factor 17 | 78 |
| *Tead1* | TEA Domain Transcription Factor 1 | 41, 79 |
| *Tead2* | TEA Domain Transcription Factor 2 | 41, 79 |
| *Klf3* | Kruppel Like Factor 3 | 80, 81 |
| *Klf4* | Kruppel Like Factor 4 | 82 |
| *Sulf1* | Sulfatase 1 | 83 |
| *Wt1* | WT1 Transcription Factor | 84, 85 |
| *Ankrd1* | Ankyrin Repeat Domain 1 | 86, 87 |
| *Cd247* | CD247 Molecule | 88 |
| *Hbegf* | Heparin Binding EGF Like Growth Factor | 89 |
| *Vegfc* | Vascular Endothelial Growth Factor C | 82, 90 |
| *Nos2* | Nitric Oxide Synthase 2 | 91 |
| *Nrarp* | NOTCH Regulated Ankyrin Repeat Protein | 92 |
| *Rgs2* | Regulator of G Protein Signaling 2 | 93 |
| *Cldn5* | Claudin 5 | 94 |
| *Kit* | KIT Proto-Oncogene, Receptor Tyrosine Kinase | 95 |
| *Cd200* | CD200 Molecule | 96 |
| *Gja4* | Gap Junction Protein Alpha 4 | 97, 98 |
| *Nog* | Noggin | 99, 100 |
| *Ets1* | Ets Proto-Oncogene 1, Transcription factor | 101, 102 |
| *Hes1* | Hes Family BHLH Transcription Factor 1 | 103 |

**Supplementary Table 5C:** Reference list for endothelial and mesenchymal genes indicated in the Figure 4D (HFD vs. Chow) heat map.

| Gene | Description | Reference # |
| --- | --- | --- |
| *Apln* | Apelin | 28 |
| *Vwa1* | Von Willebrand Factor A Domain Containing 1 | 35, 36 |
| *Tuba1a* | Tubulin Alpha 1a | 43 |
| *Ezh2* | Enhancer Of Zeste 2 Polycomb Repressive Complex 2 Subunit | 86 |
| *Fsd2* | Fibronectin Type III And SPRY Domain Containing 2 | 104-106 |
| *Mest* | Mesoderm Specific Transcript | 45 |
| *Angptl4* | Angiopoietin Like 4 | 107 |
| *Me1* | Malic Enzyme 1 | 108 |
| *Cldn5* | Claudin 5 | 94 |
| *Ankrd1* | Ankyrin Repeat Domain 1 | 86, 87 |
| *E2f1* | E2F Transcription Factor 1 | 109 |
| *Hey1* | Hes Related Family BHLH Transcription Factor With YRPW Motif 1 | 64, 110 |
| *Fndc5* | Fibronectin Type III Domain Containing 5 | 111 |
| *Dsp* | Desmoplakin | 112 |
| *Trp53inp2* | Tumor Protein P53 Inducible Nuclear Protein 2 | 113 |
| *Cd36* | CD36 Molecule | 114 |
| *Hist4h4* | Histone H4 | 86, 110 |
| *Xbp1* | X-Box Binding Protein 1 | 115 |
| *Inhbb* | Inhibin Subunit Beta B | 116 |
| *Podxl* | Podocalyxin Like | 117 |
| *Egr1* | Early Growth Response 1 | 118 |
| *L1cam* | L1 Cell Adhesion Molecule | 119 |
| *Rarg* | Retinoic Acid Receptor Gamma | 120 |
| *Ace* | Angiotensin I Converting Enzyme | 121 |
| *Apoe* | Apolipoprotein E | 122 |
| *Mkl2* | Myocardin-like protein 2 | 123 |
| *Tln2* | Talin 2 | 124 |
| *Pdgfrb* | Platelet Derived Growth Factor Receptor Beta | 125 |
| *Igfbp3* | Insulin Like Growth Factor Binding Protein 3 | 126 |

**Supplementary Table 5D:**
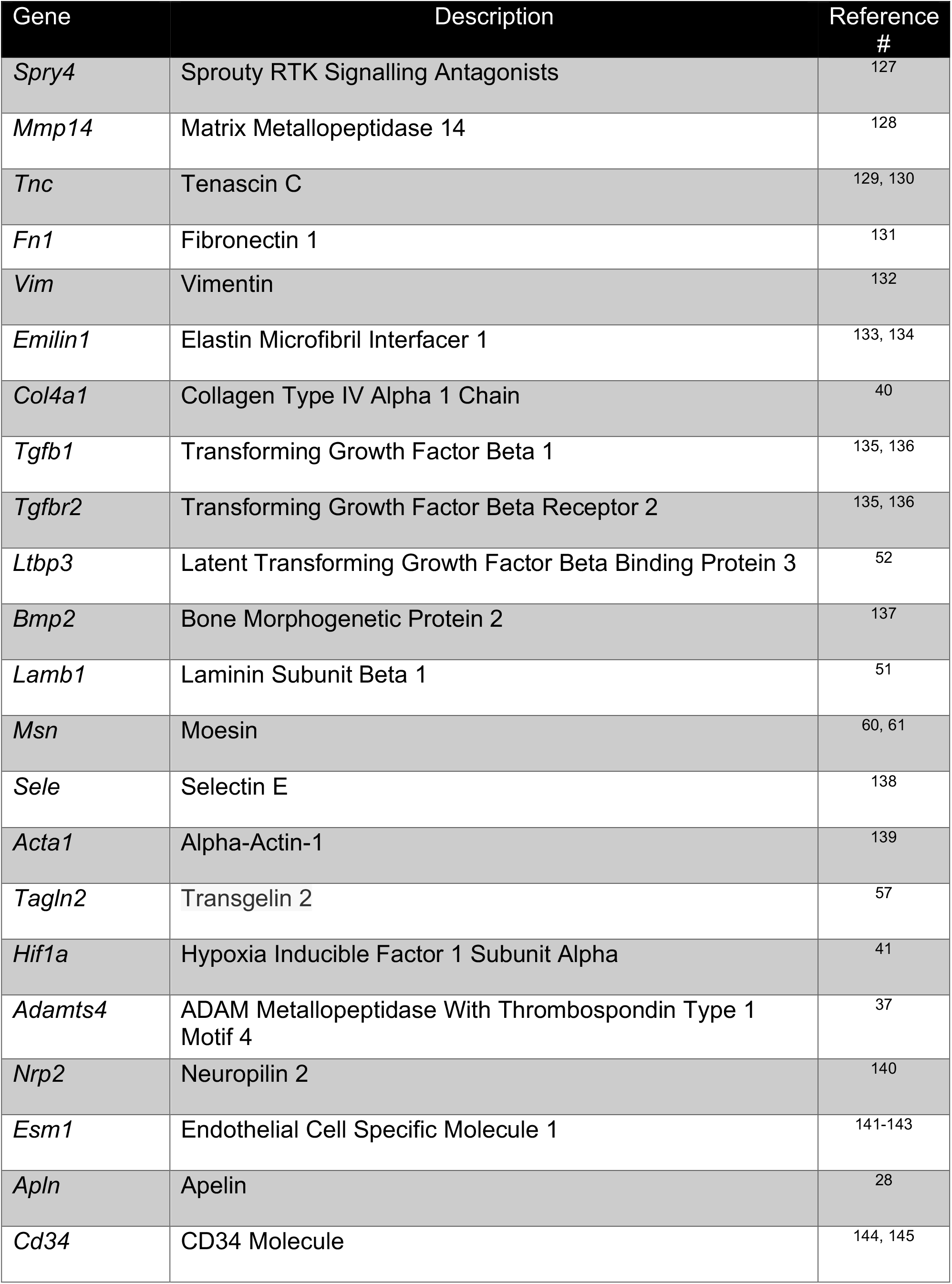
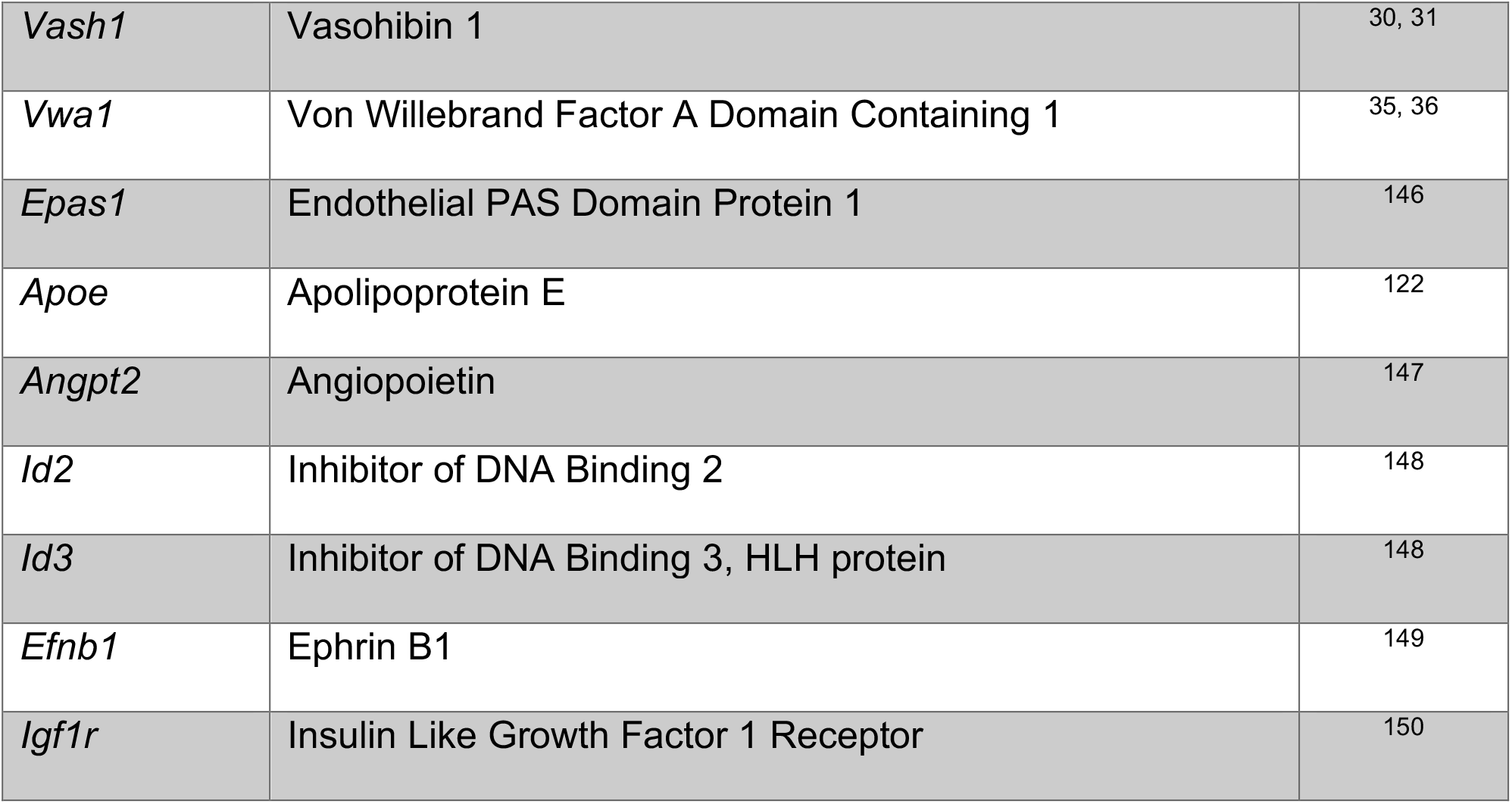
Reference list for endothelial and mesenchymal genes indicated in the Figure 4E (TAC (2) vs. Sham) heat map.

**Supplementary Table 5E:** Reference list for endothelial and mesenchymal genes indicated in the Figure 4F (TAC (7) vs. Sham) heat map.

| Gene | Description | Reference # |
| --- | --- | --- |
| *Spry4* | Sprouty RTK Signaling Antagonist 4 | 127 |
| *Esm1* | Endothelial Cell Specific Molecule 1 | 141-143 |
| *Epha2* | EPH Receptor A2 | 149 |
| *Socs5* | Suppressor Of Cytokine Signaling 5 | 62 |
| *Sele* | Selectin E | 138 |
| *Acta1* | Alpha-Actin-1 | 139 |
| *Nostrin* | Nitric Oxide Synthase Trafficking | 151 |
| *Esam* | Endothelial Cell Adhesion Molecule | 152 |
| *Cd34* | CD34 molecule | 144, 145 |
| *Tgfb1* | Transforming Growth Factor Beta 1 | 135, 136 |
| *Tnfaip2* | TNF Alpha Induced Protein 2 | 153 |
| *Ankrd1* | Ankyrin Repeat Domain 1 | 86, 87 |
| *Mmp14* | Matrix Metalloproteinase 14 | 128 |
| *Vwf* | Von Willebrand Factor | 154 |
| *Col4a1* | Collagen Type IV Alpha 1 Chain | 39, 40 |
| *Tnc* | Tenascin C | 129, 130 |
| *Col4a2* | Collagen Type IV Alpha 2 Chain | 39, 40 |
| *Stat3* | Signal Transducer And Activator Of Transcription 3 | 155 |
| *Ecm1* | Extracellular Matrix Protein 1 | 156 |
| *Zeb1* | Zinc Finger E-Box Binding Homeobox 1 | 155 |
| *Emilin1* | Elastin Microfibril Interfacer 1 | 133, 134 |
| *Sparc* | Secreted Protein Acidic And Cysteine Rich | 38 |
| *Dsp* | Desmoplakin | 112 |
| *Notch3* | NOTCH Receptor 3 | 125, 157 |
| *Pdgfrb* | Platelet Derived Growth Factor Receptor Beta | 125 |
| *Epha4* | EPH Receptor 4 | 149 |
| *Ddr2* | Discoidin Domain Receptor Tyrosine Kinase 2 | 158 |
| *Zeb2* | Zinc Finger E-Box Binding Homeobox 2 | 82 |
| *ApoE* | Apolipoprotein E | 122 |
| *Sox7* | SRY-Box Transcription Factor 7 | 159 |
| *Cxcl9* | C-X-C Motif Chemokine Ligand 9 | 160 |

